# ENCODING OF LIMB STATE BY SINGLE NEURONS IN THE CUNEATE NUCLEUS OF AWAKE MONKEYS

**DOI:** 10.1101/2021.03.17.435880

**Authors:** Christopher Versteeg, Joshua M. Rosenow, Sliman J. Bensmaia, Lee E. Miller

**Affiliations:** Northwestern University; University of Chicago; McCormick School of Engineering, Northwestern University

**Keywords:** Proprioception, Sensory Gain Modulation, Cuneate Nucleus, Monkey, Reaching, Single neurons

## Abstract

The cuneate nucleus (CN) is among the first sites along the neuraxis where proprioceptive signals can be integrated, transformed, and modulated. The objective of the study was to characterize the proprioceptive representations in CN. To this end, we recorded from single CN neurons in three monkeys during active reaching and passive limb perturbation. We found that many neurons exhibited responses that were tuned approximately sinusoidally to limb movement direction, as has been found for other sensorimotor neurons. The distribution of their preferred directions (PDs) was highly non-uniform and resembled that of muscle spindles within individual muscles, suggesting that CN neurons typically receive inputs from only a single muscle. We also found that the responses of proprioceptive CN neurons tended to be modestly amplified during active reaching movements compared to passive limb perturbations, in contrast to cutaneous CN neurons whose responses were not systematically different in the active and passive conditions. Somatosensory signals thus seem to be subject to a “spotlighting” of relevant sensory information rather than uniform suppression as has been suggested previously.

## Introduction

Proprioception plays a critical role in our ability to move, as demonstrated by the severe deficits that occur when it is absent (Proske & Gandevia, 2012; Sainburg, Ghilardi, Poizner, & Ghez, 1995). In the periphery, proprioception relies on several classes of mechanoreceptors. While joint receptors and Golgi tendon organs also contribute, muscle spindles are the primary receptor underlying proprioception (Proske & Gandevia, 2012). Because each spindle signals stretch of the muscle within which it is embedded, responses vary with movement direction, peaking for movements that lead to the greatest stretch. This characteristic may give rise to the sinusoidal tuning curves that have been described in somatosensory cortex (London & Miller, 2013; Prud’homme & Kalaska, 1994). A major challenge in studying proprioception is that both spindle sensitivity and signal transmission through the cuneate nucleus are modulated by descending inputs (Dimitriou, 2014; Ghez & Pisa, 1972) so proprioceptive responses are liable to differ for actively generated and passively imposed limb movements.

In the present study, we sought to characterize the proprioceptive response properties of CN neurons in the context of arm movements. First, we examined the degree to which CN neurons are tuned to reach direction. Observed patterns of spatial tuning suggest that individual CN neurons receive convergent input from one or only a few muscles. Second, we investigated whether CN responses to kinematically similar movements depend on whether they are produced actively or imposed on the limb. We found that the responses of proprioceptive neurons were typically potentiated during active movement but this systematic potentiation was not observed in cutaneous neurons. We speculate about why these two streams of somatosensory information may be modulated differently during active movements.

## Methods

All surgical and experimental procedures were fully consistent with the guide for the care and use of laboratory animals and approved by the institutional animal care and use committee of Northwestern University under protocol #IS00000367.

### Behavioral task

We trained three monkeys to perform a modified center-out (CO) reaching task. Each monkey grasped a handle attached to two-link manipulandum constrained to a horizontal plane. The monkeys used the position of the handle to control a cursor displayed on a vertical screen. Each trial began when the monkey moved the cursor to a target at the center of the screen. After a random delay of 0.5-1.2 seconds, an outer target appeared in one of eight locations spaced equally on a circle at a distance ranging from eight to 12 cm from the center target depending on the monkey (example trajectories in fig 1A). Some experimental sessions with monkey Bu had only four targets, one along each of the cardinal directions. Following a tone cue and the disappearance of the center target, the monkey had two seconds to reach to the outer target and hold it for a random interval between 0.1 and 0.2 seconds. If the monkey correctly performed these steps, it received a liquid reward.

**Fig 1:**
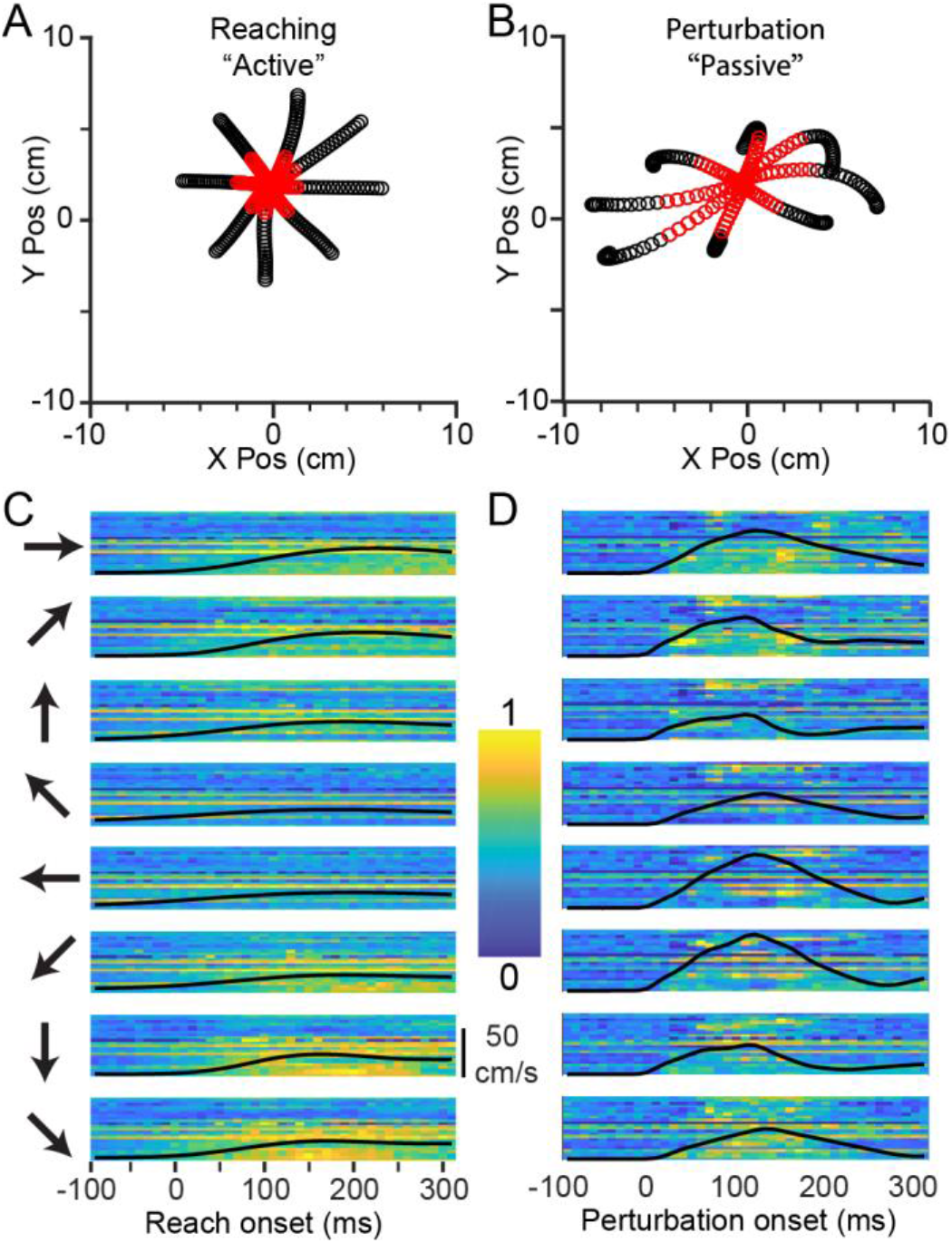
CN activity during Center-Out reaching and limb perturbation. *A:* X/Y plot of mean handle position during reaching (−100 ms to +300 ms from movement onset), averaged across ~120 trials per direction for monkey Sn. Red symbols are in the window from 0 ms to 130 ms. *B:* Corresponding plot during perturbation trials. Significant asymmetries can be seen due to the non-uniform impedance of the hand. *C:* Neural firing rates during reaching in 8 directions, indicated by the arrows to the left of the plots. Each row of pixels represents a single CN neuron, with color indicating the normalized firing rate. The black line superimposed on the image is the speed of the hand, normalized to the fastest hand speed in either the active or passive condition. *D:* Firing rates during passive trials, as in panel C.

On some trials, we imposed a force perturbation (either 2.0 or 2.5N, depending on the size of the monkey) during the center-hold period which pushed the hand in one of the eight target directions with kinematics that roughly matched that of the initial, active reaches (Fig 1A). The robot delivered the force for 125 ms, begun prior to the appearance of the outer target, but after the monkey had been holding for at least 0.3 seconds (examples in Fig 1B). In the passive trials, we confined our analyses of the neural responses to a 130 ms window beginning at bump onset to exclude the potential reafferent input due to voluntary movement. We analyzed active reaches tor 400 ms beyond movement onset, unless otherwise noted. To determine movement onset for active reaches, we found the time between the go cue and the end of the trial at which the handle acceleration crossed half its maximum, then walked backwards until we found a hand speed minimum.

### Data collection

We implanted 96-channel iridium-oxide arrays (Blackrock Microsystems) with an electrode length of 1.5 mm in three monkeys. We targeted all implants for the right CN, which receives inputs from the right arm. Detailed surgical procedures have been described previously (Suresh et al., 2017). For monkey Bu, we used a standard 10×10 shank array. For two subsequent monkeys, we maximized the area of CN sampled by implanting 8×12 shank rectangular arrays, thereby avoiding most of the gracile and trigeminal nuclei which lie medial and lateral to CN, respectively (Fig 2A). Receptive field mappings revealed areas of each array receiving inputs from the lower limb (gracile) and face (trigeminal). We used this somatotopic organization, which was conserved across time, to eliminate from consideration neurons with receptive fields not on the upper limb and torso.

**Fig 2:**
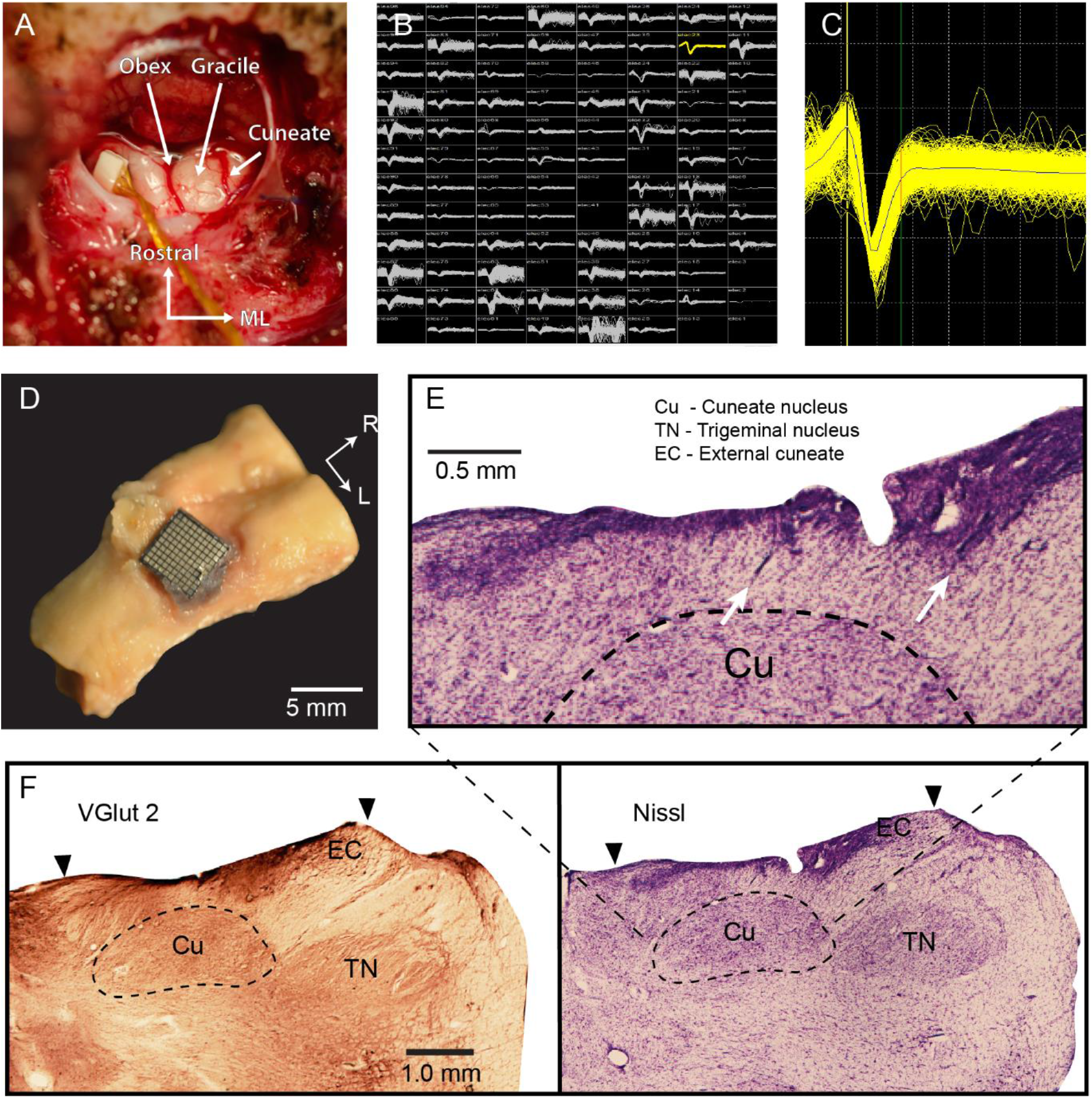
Electrode arrays implanted in dorsal brainstem yield single neuron recordings from the cuneate nucleus: *A:* Intraoperative exposure of the dorsal brainstem and cuneate nucleus following implantation of a Floating Microelectrode Array (Microprobes for Life Sciences) in an early monkey, not used in this study. The obex and cerebellar tonsils are in the center of the image. Gracile nucleus is the structure immediately lateral to the midline, with the main CN further lateral. *B:* Screenshot of the recordings across an implanted Utah array (monkey Sn). *C:* An example single neuron from monkey Sn. *D:* Histological examinations of monkey La showed that the implant successfully targeted the main CN. Brainstem with the Utah array in place. *E:* Arrows mark electrode tracks leading into the main CN. *F:* Staining by Vglut2 (left) and Nissl (right) sharply delineate the boundaries of the CN and trigeminal nuclei. Main CN (Cu) begins at ~0.5 mm depth and extends to ~2 mm. External cuneate (EC) is more lateral and shallower. Trigeminal nucleus is farther lateral. Black arrows indicate the mediolateral extent of the Utah array.

We simultaneously recorded cursor position, timestamps indicating trial events, and neural data while the monkey performed the task. We bandpass filtered the neural recordings between 250 Hz and 5000 Hz, and set a voltage threshold manually on each channel to record single neuron activity in 1.6 ms snippets surrounding each threshold crossing (Fig 2B,C). We sorted the snippets in Offline Sorter (Plexon Inc.) using waveshape and interspike interval to isolate single neurons. Neurons in CN can fire spike doublets at approximately millisecond intervals. During these high-frequency bursts, the waveshape changes, causing two clearly separable clusters in Offline Sorter. Cross-correlograms between the spike times of snippets in two such clusters have a characteristic profile, with smaller of the two waveforms reliably lagging the larger waveform (supplementary fig 1). We combined all the waveforms in these pairs of clusters to avoid double counting single neurons. We placed all the sorted spikes into 10 ms wide bins and convolved the resulting counts with a 20 ms, noncausal Gaussian kernel to produce a smoothly varying firing-rate signal for subsequent analyses.

### Histology

To confirm that our implantation procedure was appropriate to target the main CN, we performed histology on one monkey (monkey La, not included in this paper due to low neuron yield; monkeys Sn and Cr are still in use in other experiments) that had a CN implant like that of monkey Bu. The monkey was deeply anesthetized and perfused with saline followed by paraformaldehyde solution. We removed the brainstem, placing it in 5% Normal Buffered Formalin (NBF) for several weeks. The tissue was then placed in 30% sucrose in 0.1M Phosphate Buffer (PB) until it sunk. The dura and microelectrode array implant were removed, and the brainstem was blocked and mounted on a freezing microtome and sectioned coronally into 50µm sections. Tissue intended for immunohistochemical processing by VGluT2 staining was placed in 0.1M Tris-Buffered Saline with 0.1% sodium azide, while tissue for Nissl staining was placed in 5% Formalin.

#### Immunohistochemistry

The brainstem tissue was rinsed 3×5min in 0.1M Phosphate-Buffered Saline (PBS), and quenched for 10 min in 3% hydrogen peroxide in PBS. All processing was performed at room temperature. Sections were rinsed 3×5min in PBS, blocked for 2 hrs in 5% horse serum with 0.05% Tritin X-100 in PBS, and incubated overnight in the primary antibody (MsαVGluT2, Millipore, 1:5000, binding specific to vGluT2 receptors (Balaram, Young, & Kaas, 2014)) diluted in blocking solution. The tissue was then rinsed 3×5min in PBS, and placed in the secondary antibody solution (HsαMs, Vector Labs, 1:500) diluted in blocking solution for 1hr 45min, rinsed 3×5min in PBS, and incubated in the avidin-biotin complex in PBS for 2 hrs. Sections were rinsed 3×5min in PBS, developed in a solution of 0.5% DAB, 0.05% nickel ammonium sulfate and hydrogen peroxide in 0.1M PB and given a final rinse in PBS. Sections were mounted on gelatin-coated slides, dried, and cover slipped with DPX.

#### Nissl Staining

Sections were mounted out of 0.1M PB onto gelatin-coated slides and left to dry overnight. The tissue was then placed in a 1:1 chloroform and ethanol solution and sent through an ascending ethanol series into xylenes for a 15min incubation. The tissue was then put through a descending ethanol series into water and placed into the Nissl substance for 15min, followed by differentiation in 70% ethanol with acetic acid, and back through the ascending ethanol series into xylenes. The sections were cover slipped out of xylenes in DPX.

### Receptive field mapping

During our experiments, we found some neurons that responded to body segments other than the proximal arm and upper torso. To exclude these neurons from our analysis of limb movement, we mapped the receptive fields (RFs) of neurons under light ketamine or dexmedetomidine sedation after all reported experimental sessions. These RF mappings typically took one to two hours, limited by sedation time and animal tolerance for manual mapping. We excluded from our analyses all neurons that had receptive fields on the forearm, hand, legs, lower torso and head or face. We also removed neurons that had a stereotypical bimodal passive tuning curve that was indicative of receptive fields on the hand (for example, see supplementary fig 2). Gross somatotopic arrangement of RFs was consistent across long time periods (fig 3), allowing us to target electrodes that reliably had proximal limb RFs.

**Figure 3:**
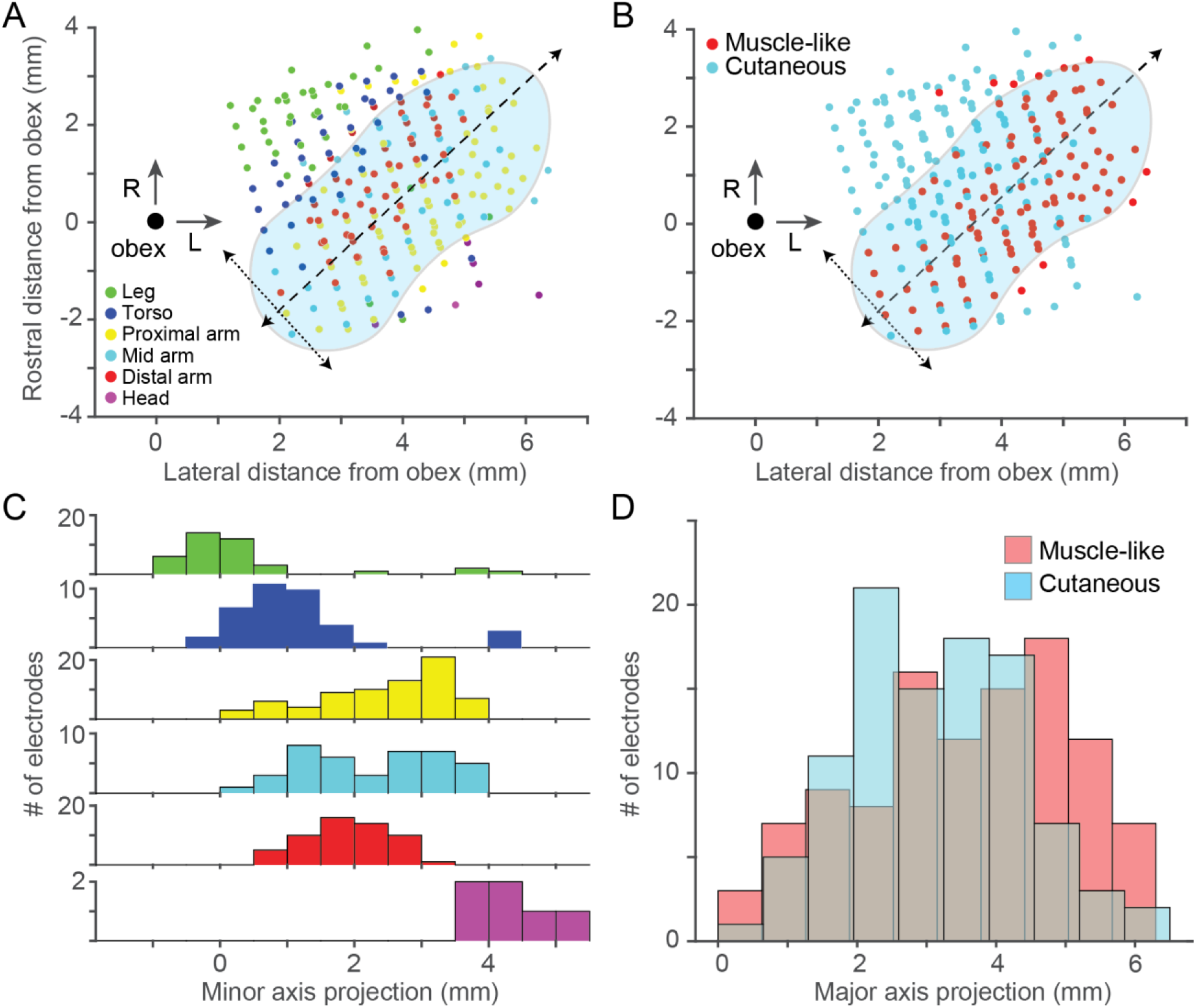
Receptive field location and modality across monkeys. A: Scatter plot of receptive field locations as a function of the location of the recording site relative to obex (large black point). Each point represents a recording site in the dorsal medulla from one monkey. Color of points denotes the most common receptive field location for a given electrode. Approximate location of CN is show in blue, with its major (dashed) and minor (dotted) CN axes overlaid. RF locations appear to vary primarily along the minor axis. “Proximal arm” included shoulder related receptive fields, “Mid arm” included RFs around the elbow, and “Distal arm” included all forearm and hand related RFs. B: Modality as a function of electrode location. As in A, symbol color indicates the most common modality. C: Histogram of receptive field location along the minor axis in A, relative to the obex. RFs progressed systematically along minor axis from lower limb (green) to head/face (purple). D: Histogram of receptive field type along the major axis. There was a weak bias for muscle-like RFs away from the obex.

To find neurons with apparent cutaneous input, we brushed the skin around the arm and torso while listening to pulses from discriminated action potentials. Cutaneous receptive fields were of highly variable size; some responded to brushing of skin over large areas, while others had focal receptive fields, often on the hand. Due to methodological limitations, we did not test for Aδ receptive fields, joint receptor afferents, or Golgi tendon organ input.

To find putative spindle afferents, we began with passive arm movements to determine articulations in which the neuronal firing rate increased and used that information to guide palpation of muscles that lengthened during those articulations. We then applied 100-Hz vibration to the belly of these muscles, using either an electrodynamic LDS V101 shaker (BRÜEL & KJÆR) or smaller vibration motors. This stimulus has been shown to activate primary muscle spindle afferents (Fallon & Macefield, 2007; Proske & Gandevia, 2012). Often, CN neurons responded only to vibration of small regions of the muscle belly.

We classified a neuron as a putative recipient of muscle spindle input (“spindle-receiving”) if it responded to the lengthening, and either vibration or palpation of a given muscle, but not to stroking of the skin overlying the muscle. When testing a putative muscle spindle-receiving neuron, we vibrated the muscle through different patches of skin (by manually displacing the lax skin) to confirm that the response was caused by vibration of the muscle and not the overlying skin. We found occasional neurons that responded to vibration of more than one muscle, typically adjacent synergist wrist flexors. Whether this was due to convergence of multiple muscle receptors onto a single CN neuron or vibration spreading to adjacent muscles is difficult to determine with certainty. We defined any neuron that consistently and selectively responded to passive movements of the limb, but with an RF that we were not able to localize to a single muscle or cutaneous field as “muscle-like”. This included the spindle-receiving neurons.

### Motion tracking

In one monkey, we used three video cameras to record the movements of the monkey’s arm. We triggered frame collection with a 30 Hz pulse transmitted from our data collection system, simultaneously recorded as an analog input for post-hoc alignment of neural, task, and video data. We used a publicly available package (DeepLabCut (Mathis et al., 2018)) to infer 10 locations on the monkey’s arm after training on ~200 hand-labelled reference images. We reconstructed 3D coordinates of each location based on four separate camera views. Based on the output from DeepLabCut, we used Opensim (Delp et al., 2007) and a 3D musculoskeletal model of a macaque arm with 7 degrees-of-freedom (Chan & Moran, 2006) to compute the lengths and velocities of 39 muscles. We binned these data at 10 ms and aligned them in time with the neural data.

### Spatial tuning curves and preferred directions

We calculated the mean firing rate and its 95% confidence interval for each neuron across trials in a 130 ms period beginning at perturbation onset, or in active trials, the 200 ms surrounding the peak hand speed in each direction. In addition to the classic method of fitting a sinusoid to trial-averaged data (Georgopoulos, Kalaska, Caminiti, & Massey, 1982; Prud’homme & Kalaska, 1994), our lab has begun to compute preferred directions (PDs) by fitting models from hand velocity to the smoothed firing rates of each neuron using Poisson Generalized Linear Models (GLMs) (Chowdhury, Glaser, & Miller, 2020). This latter approach is sensitive to variability in reach kinematics across trials and can be applied to random-target reaching tasks as well as center-out tasks. Here, we concatenated all trials and placed the data is 50 ms bins. In Eq 1, λ and α represent the time-varying and baseline firing rates (spikes/sec), respectively, of a given neuron. β is the weight vector for the x and y components of velocity. We computed a PD from the GLM by taking the inverse tangent of the ratio of the y and x velocity weight vectors, β. We used bootstrap sampling across data points to generate 95% confidence intervals on the PD.

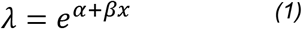

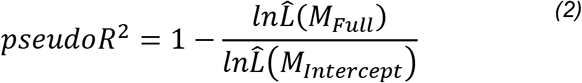

### Neural tuning metrics

We classified neurons as “Active Tuned” if there were statistically significant differences in an F-test across reaching directions with a cutoff of p < 0.01. Similarly, neurons were “Passive Tuned” if they met the same criterion for bump-evoked responses. Neurons could be “Active Tuned”, “Passive Tuned”, or both. We considered neurons “Sinusoidally Tuned” if the PD confidence interval had a total width of 90 degrees or less. Neurons could be “Active Sinusoidally Tuned”, “Passive Sinusoidally Tuned” or both.

We found the time a neuron modulated relative to movement onset for each target direction (supplementary fig 3) by computing the trial-averaged firing rate in 10 ms bins from 100 ms prior, to 200 ms after movement onset. We found the first time at which this average rate was outside the 99.9 percentile of the baseline firing rate (from 150 ms to 100 ms prior to movement onset) for two consecutive bins. We computed the latency for passive movements in a similar manner, using a baseline window from 100 to 50 ms prior to perturbation onset, testing for changes from 50 ms prior to the bump to 100 ms after the bump.

### Analysis of simulated spindle-receiving CN neurons

To determine the extent to which the representation of movement direction within CN resembles that of the periphery, we compared the spatial tuning of CN neurons to that expected from their apparent muscle spindle inputs. This process had several steps. We computed typical length changes of arm muscles while a monkey performed the CO task using motion tracking data from a single session of monkey Sn. We simulated spindle firing rates by passing the lengthening velocity of each muscle through a power law with coefficient of 0.5 (Houk, Rymer, & Crago, 1981). We set firing rates during muscle shortening to zero. We scaled each spindle output to a firing rate of 50 Hz at near-maximal lengthening speed (90^th^ percentile). Treating this rate as the time-varying λ of a Poisson distribution, we sampled randomly to generate firing rates for each simulated spindle on each trial. Finally, we used a linear model to determine PDs for the simulated spindles from the velocity of the hand as we did for CN neurons.

We computed the PDs for simulated CN neurons that each received input from a single randomly chosen muscle spindle from muscles distributed throughout the proximal arm. The number of muscle spindles in each muscle is roughly proportional to the square root of the muscle’s mass (Banks & Stacey, 1988). We estimated the mass of each muscle by the multiplying the pulling force (proportional to cross-sectional area) by the length of the muscle, both of which were included in our musculoskeletal model. Thus, we assumed that the number of muscle spindles in each muscle was proportional to the square root of its pulling force times the length of the muscle. We simulated 1000 muscle spindle-receiving CN neurons, apportioned across the muscles on this basis. From this population, we computed PD distributions based on the kinematics for active reaches.

### Sensitivity analyses

To estimate the sensitivity of CN neurons to hand movements, we used the x and y components of hand velocity as input to linear models that predicted the smoothed firing rate of each neuron. The length of the weight vector was that neuron’s sensitivity to velocity and quantifies the expected change in firing rate for an increase of one cm/s in the direction of the neuron’s velocity PD.

Due to the anisotropy of the limb and idiosyncrasies of a monkey’s task performance, the perturbations did not produce kinematics perfectly matching those of the reach. If firing rates are a nonlinear function of speed, such as the power law observed in muscle spindles (Houk et al., 1981), mismatched movement speeds across conditions would bias the apparent sensitivity. To address this potential confounding factor, we matched the input velocity domains of the data used to train the models. We found separate static 2D distributions of firing rates as a function of velocity for the active and passive trials. For each reach-velocity datapoint, we found the distance to the nearest passive datapoint, in an approach analogous to a nearest neighbor method. If this distance was greater than 3 cm/s, we excluded the active point, as it had no near neighbors. We repeated this process to exclude passive data that did not have active neighbors. The result was training data in which the active and passive movements had matched velocity domains. The data windowing did not substantially alter the results of the sensitivity analyses; we demonstrate the data windowing and its effects on the results of this analysis in supplementary fig 4.

To compute whether a neuron’s movement sensitivity differed significantly between the active and passive conditions, we bootstrapped, across trials, a confidence interval on the difference between active and passive sensitivities for each neuron. If the mean of this metric was positive and the 95% confidence interval did not include zero, the neuron was more sensitive in the active condition; if the mean was negative and the 95% confidence interval did not include zero, the neuron was significantly less sensitive.

## Results

We recorded the responses of neurons with receptive fields (cutaneous or proprioceptive) on the proximal arm while the animals performed a modified Center-Out reaching task that included force pulse perturbations applied to the robot handle during the center-hold period. Unless otherwise specified, the data were obtained in two sessions with each monkey, separated by at least three weeks to reduce the likelihood of double-counted neurons.

### Somatotopic organization of CN is similar across monkeys

First, we examined the somatotopic organization of the CN by systematically mapping the receptive field types and locations across the arrays. Using intra-operative photos of array placement, we found the coordinates of each array relative to the obex. We then plotted the most common RF location (i.e., legs, trunk etc.; Fig 3A) and modality (muscle-like, cutaneous; Fig 3B) for each electrode). Receptive field locations varied systematically along the minor axis cutting through CN (dotted arrows in Fig 3A, projected onto axis in 3C). This progression reflects the transition from the gracile nucleus to CN, and finally to the trigeminal nucleus. RF locations on the arm were largely conserved along the major axis, possibly corresponding to the CN subnucleus known to receive primary inputs from distal cutaneous receptors (Loutit, Vickery, & Potas, 2021). These results are consistent with our histological results from one monkey (Fig 2F), which indicate that the array likely penetrated through the external CN, to record from rostral portions of the main CN. We could not confirm this independently for all monkeys, for which histology has not been completed. The orientation of the major and minor axes departs from strictly mediolateral because of the sharply lateral bend of the brainstem and nuclei just rostral to the obex. RF type varied along the major axis, with muscle-like response properties slightly more common farther from the obex (fig 3B,D).

### Localized vibratory stimulation robustly activates CN neurons

Having identified joints that appeared to be within the RF of a given CN neuron, we characterized the spindle input to that neuron by applying vibration to the belly of muscles that articulate that joint. Figure 4 shows the response of a CN neuron to vibration of the brachialis muscle, presumably due to the activation of its muscle spindles. As in this example, neural responses typically increased and became phase locked with the vibration. We found that many of these spindle-receiving neurons required the vibration be delivered quite precisely within a given muscle to be effective, suggesting that CN neurons may not even receive input from spindles throughout a given muscle.

**Fig 4:**
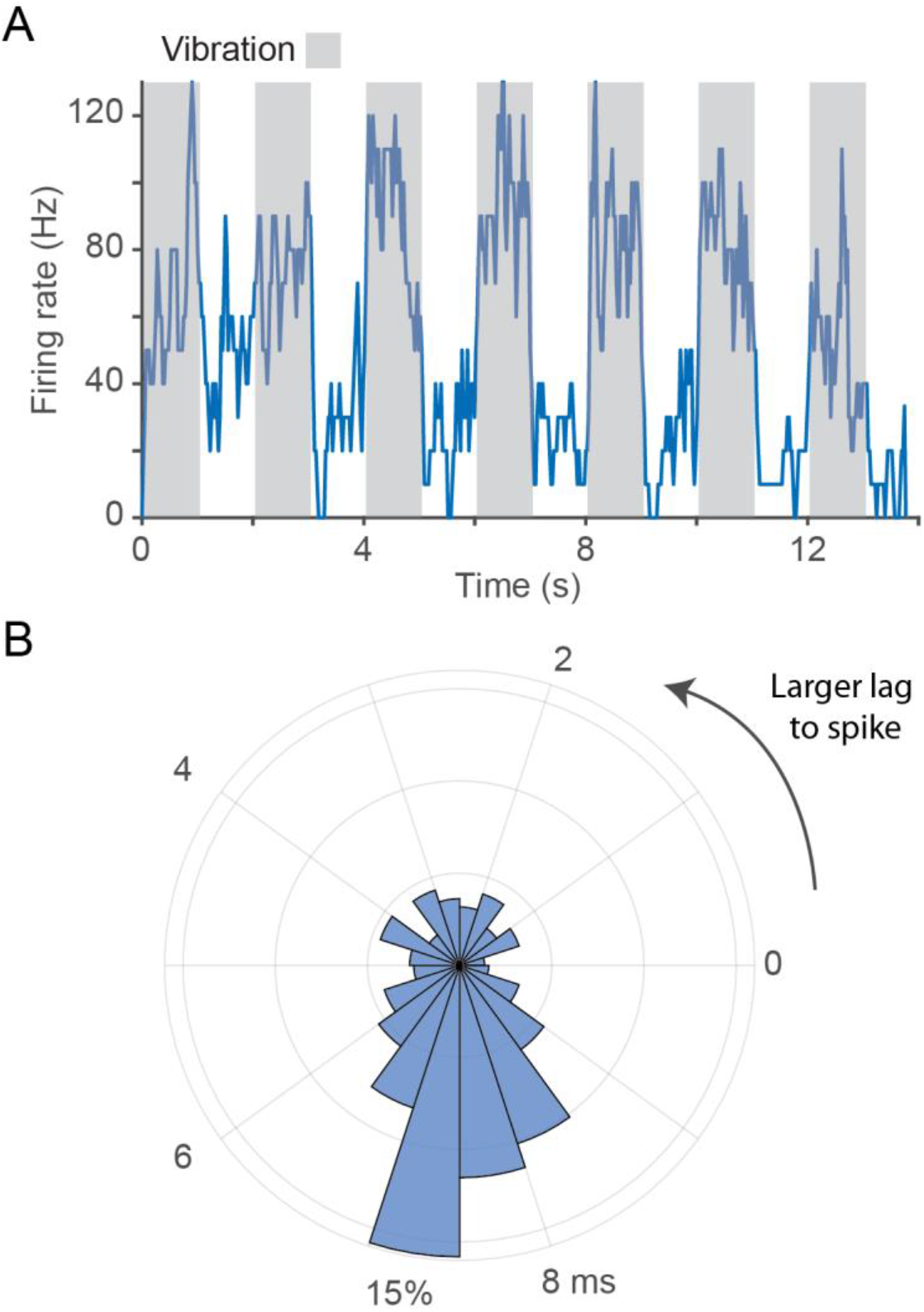
Responses to muscle vibration of a spindle receiving neuron. *A:* Response of a CN neuron to 100 vibration applied to the brachialis muscle belly. Grey regions indicate the stimulation epoch. The neuron’s firing rate rose quickly to 100 Hz and returned to baseline immediately when the vibration stopped. *B:* Phase locking between the vibration peaks and potentials. We computed a phase histogram between the peak voltage applied to the stimulation and evoked spikes. The peak at ~7.5 ms indicates that the vibrator peak led this neuron’s spikes with a reliable latency. Some of the breadth of the peak is certainly due to the sinusoidal nature of the stimulus.

Next, we examined the degree to which CN neurons receive input from multiple muscles. In most cases, CN neurons responded to passive manipulation (or vibration) of a single joint or muscle. In a few cases (<10), we found evidence that signals from multiple (typically agonist) muscles converged onto a single CN neuron. We never found neurons that responded to muscles that were not in near proximity to one another nor did we find neurons that exhibited both cutaneous and proprioceptive responses, though due to time constraints on sensory mappings, convergence may be broader than our mappings suggest.

### CN neurons are tuned to movement direction

Figure 5 shows the responses of two representative CN neurons measured during ~50 reaches in each of eight directions, a cutaneous neuron with an RF on the axilla (Fig. 5A,B) and a spindle-receiving neuron with an RF on the triceps muscle (Fig 5C,D). The firing rates of both neurons varied with movement directions, peaking for a single target direction, with similar tuning during reaching and passive limb displacement.

**Fig 5:**
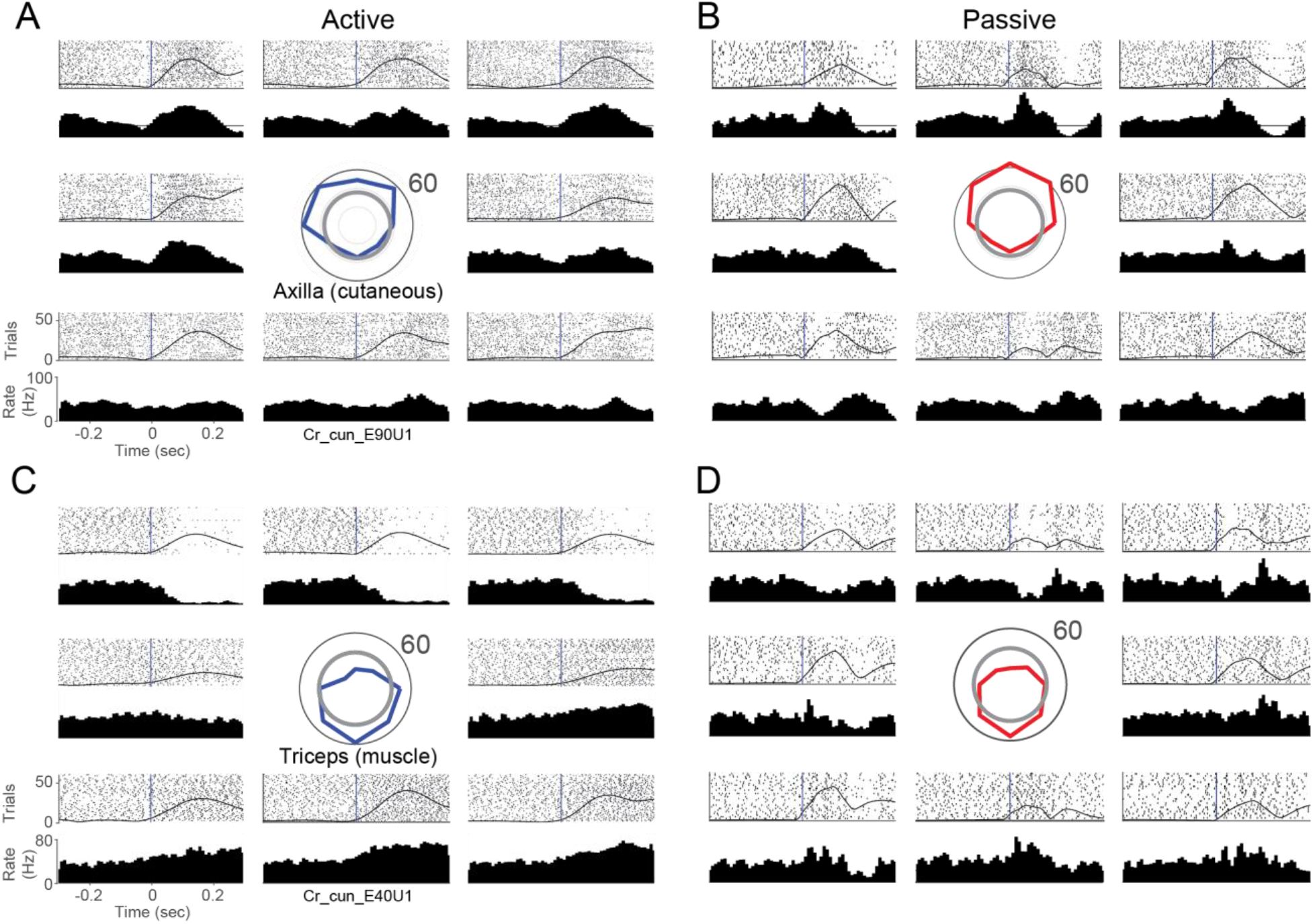
CN neurons respond robustly to active and passive arm movements: A: Responses of a CN neuron during active reaches in eight directions. RF mapping revealed that the neuron received input from cutaneous receptors in the axilla. The tuning curve (centered, blue) indicates the firing rate averaged across the 130 ms after movement onset in each direction. The grey circle illustrates the baseline rate before movement. Rasters and histograms are positioned relative to the tuning curve, to correspond to the direction of movement. The black vertical lines indicate movement onset. The hand speed is represented as a solid black line imposed over the rasters. B: Same neuron as A, for passively evoked arm movements. Passive tuning curve plotted in red at center. C,D: A second neuron, presented as in A, B, that appeared to receive input from receptors in the triceps muscle spindle.

During active reaching movements, trial-averaged firing rates of muscle-like neurons in CN were generally well fit by a cosine tuning model (Georgopoulos et al., 1982; Prud’homme & Kalaska, 1994), with average fits of r = 0.76. Cutaneous neurons yielded, on average, a cosine fit of 0.62, which was not statistically different from the muscle-like population (t-test p-value ≈ 0.10). These values are very similar to those reported previously for neurons in motor and somatosensory cortices (Georgopoulos et al., 1982; Prud’homme & Kalaska, 1994) (See supplemental fig 5 and supplemental Table 1). For compiled firing rate, sensitivity, and latency metrics, see supplemental fig 6.

Other neurons exhibited idiosyncratic responses, including unexpected dynamics at movement onset, (supplementary fig 7), potential GTO inputs (supplementary fig 8), cutaneous responses from the hand (supplementary fig 2) and forearm (supplementary fig 9).

### Distribution of CN PDs can be predicted from single-muscle receptor inputs

Next, we examined the distribution of PDs across the population of CN neurons and found it to be highly non-uniform (fig 6A,B): A large proportion of PDs fell within a single lobe pointed toward the body (near −90°) in both the active and passive conditions. This observation was consistent across monkeys (supplemental fig 10). To shed light on this result, we simulated a population of CN neurons, each with spindle input from a single muscle, inspired by the very limited convergence we found for vibration-evoked responses in muscle-like neurons (see Methods). The resulting distribution of simulated PDs featured a mode at −90°, much like that that of the CN neurons, but also another mode at 90° (Fig 6C).

**Figure 6:**
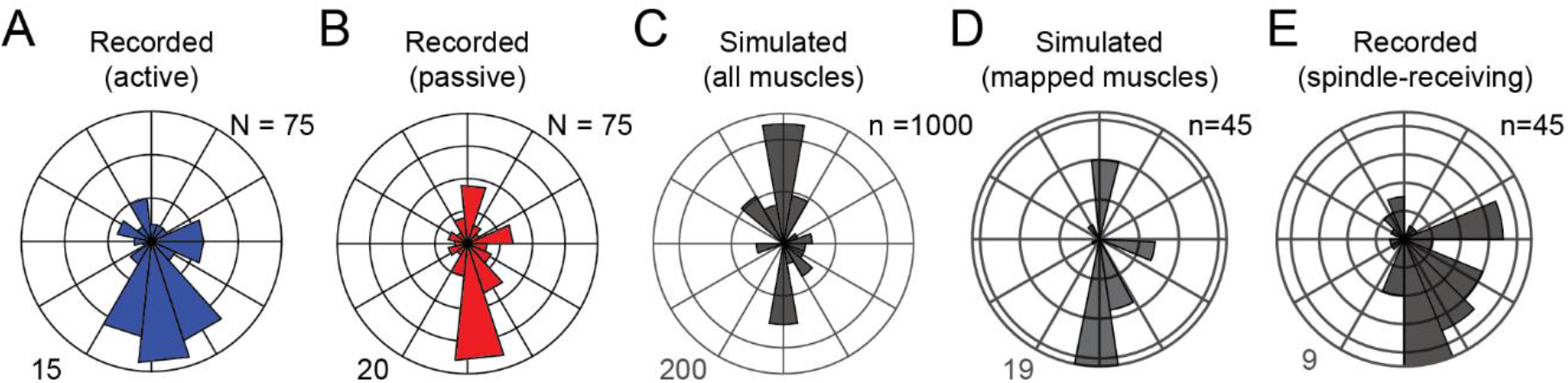
Preferred direction distributions for simulated and actual CN neurons. A) Polar histogram of active PDs combined across monkeys (N= 75 neurons). Outer circle represents 15 neurons with PDs in that bin. All subsequent plots in this figure have the same layout as A. Neurons included in this figure were sinusoidally tuned in both active and passive conditions, from CN regions of the array, and appeared to receive inputs from the upper trunk, shoulder or proximal arm. B) Passive PD distribution for CN neurons. C) PD distribution for all 1000 simulated CN neurons receiving input from a muscle spindle of a single randomly chosen muscle in the proximal arm. D) PD distribution for simulated CN neurons, having inputs corresponding to those actually mapped for recorded neurons. E) Actual PD distribution of the same spindle-receiving neurons in D.

This strongly bimodal distribution of simulated spindle-driven neurons reflects the biomechanical non-uniformity of muscles, which predominantly drive arm movements toward and away from the body. A consequence of this anisotropy in muscle pulling directions is that we can push and pull objects with greater strength than we can move them from side to side. The lack of neurons with PDs pointing away from the body suggests that we recorded neurons with a somatotopically biased set of RFs, namely a preponderance of neurons driven by lengthening of elbow extensors and shoulder flexors and lacking neurons driven by their complements. When we limited the inputs to our simulated neurons based on the mapped RFs of our recorded CN neurons, the two PD distributions matched more closely (fig 6D,E). Even at the single-neuron level there was a reasonable correspondence between the PD of the recorded neurons and their modeled counterparts. While prediction accuracy was poorer for CN neurons that received inputs from muscles in the back (which tend to be multi-layered, broad, and biomechanically dissimilar), accuracy for CN neurons that received inputs from the arm was high (supplementary fig 11). These results are consistent with the view that CN neurons receive input primarily from individual muscles.

### Directional tuning of active and passive responses are similar

Next, we examined the directional tuning during actively generated movements and compared it to directional tuning during imposed limb perturbations, focusing on neurons that exhibited sinusoidal directional tuning. First, we found that the depth of modulation was correlated across conditions: Neurons that were strongly tuned in the active condition were also tuned in the passive one (fig 7A). Second, we found that PDs were typically consistent across the two conditions (Figure 7B), with more than 50% of neurons exhibiting active and passive PDs that differed by less than 30º (Figure 7C). From these data, we conclude that CN neurons convey information about direction that is largely consistent regardless of whether limb movements are generated actively or imposed.

**Fig 7:**
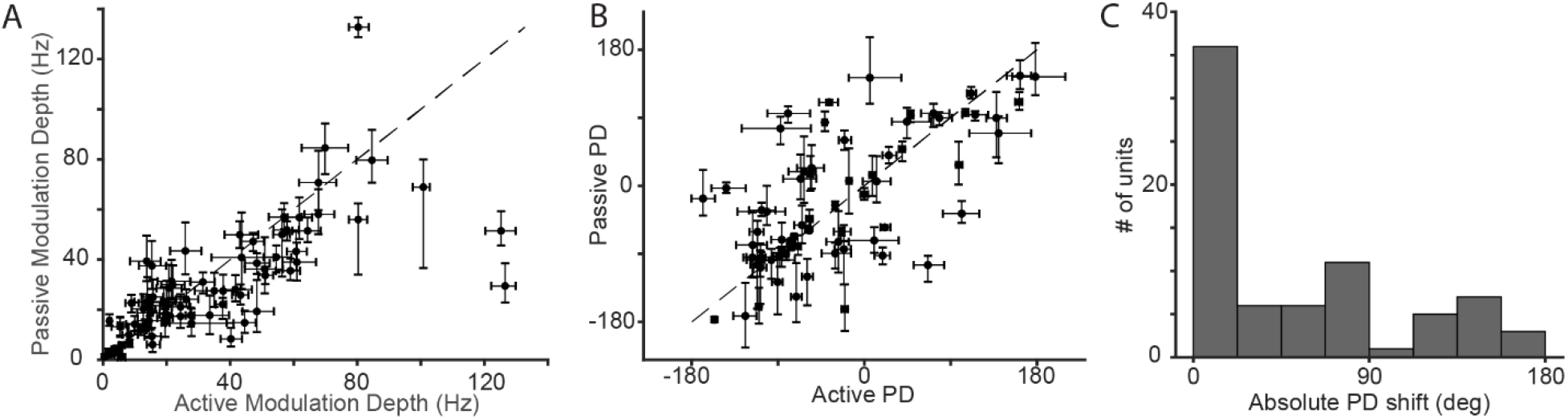
CN neurons have similar active and passive tuning: *A:* Each point represents the modulation depth of a neuron in the passive condition plotted against its active modulation depth. Error bars denote the bootstrapped 95% confidence interval of the modulation depth computed across trials. Neurons in the figure have the same inclusion criteria as those of Fig 6A. B: Each point represents the active and passive tuning direction for single proximal limb CN neurons that were sinusoidally tuned in both conditions. The black dashed line is the unity line. The error bars denote the bootstrapped 95% confidence interval on the PD. *C*: Histogram of the absolute angle between active PDs and passive PDs.

### Response strength differs in the active and passive conditions

CN responses to tactile stimulation have been shown to be suppressed during movement (Ghez & Pisa, 1972; He, Suresh, Versteeg, Rosenow, & Bensmaia, 2019), a phenomenon that likely accounts in part for the documented decrease in cutaneous sensitivity during movement (Williams & Chapman, 2000, 2002; Williams, Shenasa, & Chapman, 1998). With this in mind, we examined the degree to which such a gating phenomenon occurred in our sample of CN responses. Specifically, we compared the strength of the response evoked in CN neurons in the active vs. passive movement conditions. As the kinematics were not identical in the two conditions, we selected a subsets of trials with similar movement kinematics. Furthermore, we focused the analysis on the responses of 70 neurons whose responses were sinusoidally tuned for at least one of the two conditions, 56 of which were muscle-like and the rest cutaneous.

Of the muscle-like neurons, ~40% (22) were potentiated during active reaching and 14% (8) were attenuated; the remaining 46% produced responses that did not differ significantly in the two conditions (Fig 8A). Among the muscle-like neurons, the results were similar whether or not they were spindle-receiving. Of 14 cutaneous neurons with RFs on the upper torso and proximal arm, 28% (4) were significantly potentiated, another 28% were attenuated, and the remainder were not significantly affected. To quantify the degree of potentiation or attenuation, we projected the responses onto a “potentiation axis” orthogonal to the unity line on fig 8B. Positive projections indicate potentiation, while negative projections indicate attenuation. As a population, muscle-like neurons were potentiated during reaching, while cutaneous neurons were not (Fig 8C; two-sided t-test p < 0.05 for proprioceptive neurons, p > 0.70 for cutaneous neurons). We found that spindle-receiving neurons as well as the more general class of muscle-like neurons were similarly potentiated. We also examined the consistency of the potentiation, which varied considerably across neurons for all monkeys and found the potentiation was quite consistent across the first and second halves of experimental sessions (figure 8D). Both the sign and magnitude of the potentiation were well preserved for virtually all neurons, cutaneous as well as muscle-like.

**Fig 8:**
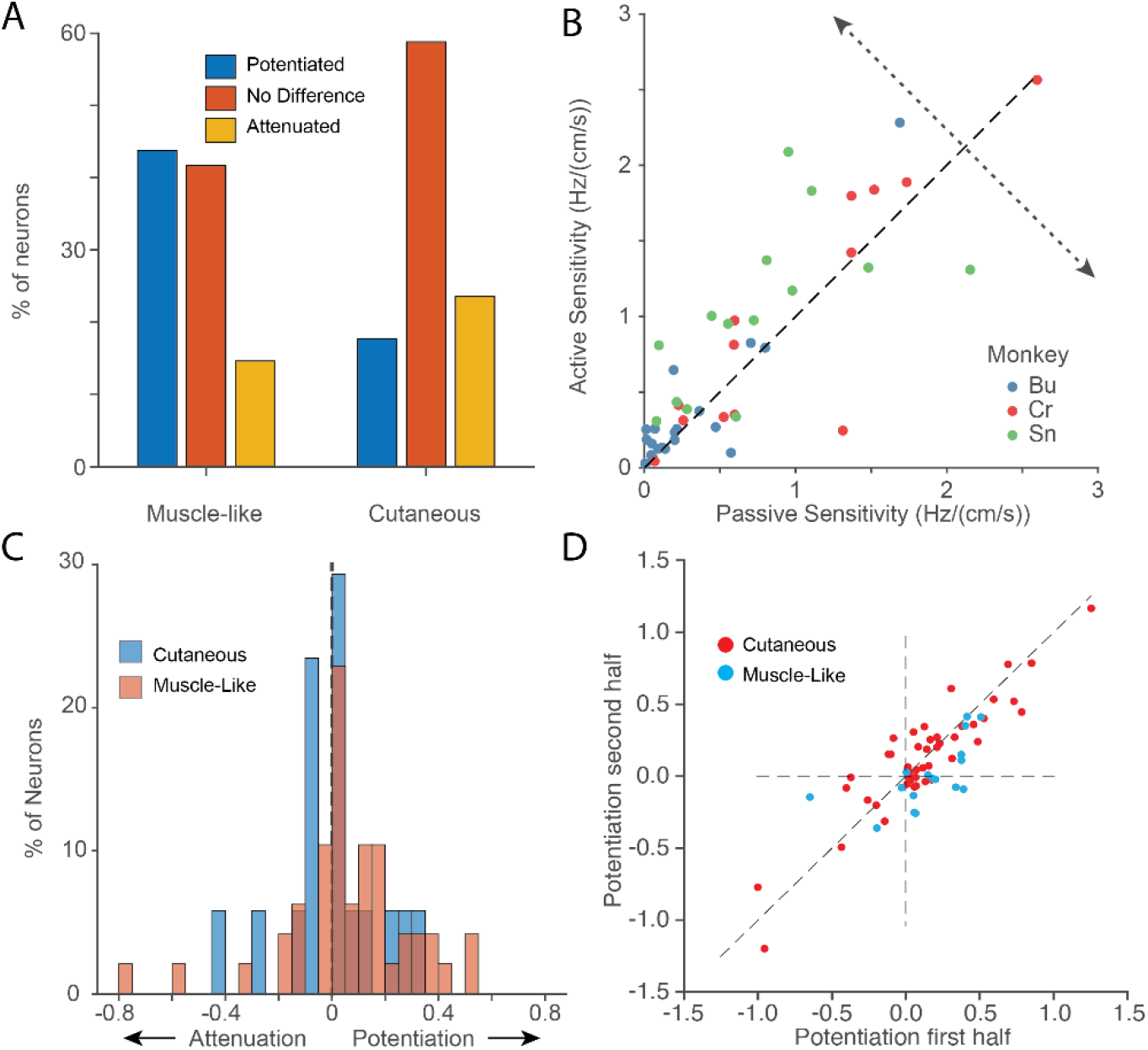
Sensitivity of neurons is modulated by movement context: A: Distribution of sensitivity changes with reaching across all neurons by modality (across two experimental sessions from each monkey). B: Scatter plot of active sensitivity as a function of passive sensitivity for all muscle-like CN neurons that were sinusoidally tuned in either condition with RFs that didn’t include the distal arm. The potentiation axis (dotted line) indicates change in sensitivity of active reaching vs. passive perturbation. Symbol color indicates the monkey from which the neuron was recorded. C: Magnitude of the potentiation across neurons. While muscle-like neurons (red) yielded positive gains, cutaneous neurons were not more significantly more prone to potentiation or attenuation (blue). Overlap between these distributions appears dark red. D: Scatter plot of the potentiation effect in the second half of a given experimental session plotted against that in the first half, for all monkeys.

## Discussion

In this study, we examined the representation of arm movements – actively generated and passively imposed – in the CN of three monkeys. First, we found that CN neurons are strongly activated during both types of movement, typically with sinusoidal directional tuning that is largely conserved between the two conditions. Second, our inability to drive CN neurons with vibrations applied to more than one muscle, and the similarity of actual CN preferred directions to those derived from the simulated spindle responses of single muscles, suggest that most CN neurons receive input from a single muscle. Third, while directional tuning is similar in the active and passive conditions for muscle-like CN neurons, their sensitivity to movement is potentiated during active reaching. This potentiation is not observed in cutaneous neurons.

### Convergence of multiple muscles onto CN neurons is limited

We never observed cross-modal convergence and found only infrequent convergence from multiple muscles. Those few neurons that appeared to have multi-muscle RFs received inputs from multiple forearm muscles. It may be that forearm muscles have higher levels of convergence than other muscle groups. It is possible that this finding reflects greater mechanical coupling between the parallel forearm muscles (Hummelsheim & Wiesendanger, 1985), but the precise placement of the vibrator, even within a single muscle, required to evoke firing argues against this interpretation.

Prior studies investigating whether afferent signals from multiple muscles converge onto individual CN neurons have yielded contradictory results. One study found that CN neurons typically respond to stretch of only one forearm muscle (Hummelsheim & Wiesendanger, 1985), with only about 25% of neurons exhibiting convergence from another muscle. In contrast, another study found that 87% of CN neurons could be excited by electrical stimulation of more than one peripheral nerve. A high percentage responded even to stimulation of both superficial and deep radial nerves (purely tactile and proprioceptive, respectively) suggesting cross modal in addition to cross-muscle convergence (Witham & Baker, 2011). A more recent study helps to reconcile these findings; Bengtsson et al. found that while CN neurons often receive input from a large number of afferents, only a small number of them strongly activate CN; the majority are “silent synapses” (Bengtsson, Brasselet, Johansson, Arleo, & Jörntell, 2013). The high levels of convergence observed with peripheral nerve stimulation may result from nonphysiological levels of synchronous inputs.

This evidence of limited convergence onto CN is supported by our ability to predict the PDs of individual spindle-receiving neurons based on the single dominant muscle in their receptive field. This was true both at the single-neuron level (primarily for CN neurons that received inputs from the arm; supplementary fig 11) as well as the population level, with one caveat. While the major node of the CN PD distribution pointing toward the body closely matched that of the simulated distribution. The latter had an additional prominent lobe pointing away from the body, which was only weakly represented in the CN distribution. This bimodal PD distribution was predicted previously for both muscle spindles (Sandbrink et al., 2020) and neurons in primary motor cortex (Lillicrap & Scott, 2013). The discrepancy between simulated and actual CN PD distributions may be explained by a sampling bias introduced by the fixed depth of the recording electrodes. Consistent with this idea, somatotopic organization in DCN-complex nuclei has been observed not only along the mediolateral and rostro caudal axes but also in depth (Loutit et al., 2021; Suresh et al., 2017). Previous investigations have found proprioceptive CN neurons over 3.5 mm deep compared to our 1.5 mm, suggesting that we may be sampling less than half of the depth-extent of CN with this array design.

For the most part, active and passive PDs were similar for CN neurons, with more than 50% differing by less than 30°. There were occasional discrepancies, which likely arise from a combination of factors including PD estimation uncertainty (Stevenson et al., 2011), altered descending drive (including gamma drive, sup. Fig 7) or convergence from unmodeled receptors, such as GTOs (sup. Fig 8).

### Modulation of CN response sensitivity during active and passive arm movements

Tactile perceptual sensitivity is attenuated during self-generated movement (Juravle, Binsted, & Spence, 2017; Schmidt, Schady, & Torebjörk, 1990). Consistent with this observation, the magnitude of evoked potentials in somatosensory cortex is also reduced during reaching (Rushton, Rothwell, & Craggs, 1981). This attenuation has been shown to occur at least in part at the level of CN, where experiments in cats showed that CN output is attenuated both by stimulation of the motor cortices (Andersen, Eccles, Oshima, & Schmidt, 1964) and during active stepping movements (Ghez & Pisa, 1972). These effects are at least partially mediated by presynaptic inhibition in the cuneate (Andersen, Eccles, Schmidt, & Yokota, 1964). These observations led to the hypothesis that afferent signals might be attenuated to reduce sensory noise, particularly during rapid, ballistic movements intended to be executed without feedback (Cohen & Starr, 1987; Rushton et al., 1981). However, more recent studies reveal a more nuanced picture: particular CN responses are potentiated when stimulation is applied to a cortical site with an RF that matches that of CN and attenuated when the RFs do not match (Palmeri, Bellomo, Giuffrida, & Sapienza, 1999). We found that about 40% of all CN muscle-like neurons were potentiated in the active condition, while only 15% were attenuated (though some quite markedly).

The responses of cutaneous nerve fibers have been shown to carry limb-kinematic information comparable to that of muscle spindles (Edin, 1992). Furthermore, activation of cutaneous afferents in a manner that mimics that occurring during arm movement biases the conscious perception of hand location (Collins, Refshauge, Todd, & Gandevia, 2005; Edin & Johansson, 1995). To the extent that cutaneous signals complement muscle-derived ones to support proprioception, one might expect that cutaneous signals would also be potentiated during active movements. In our experiments, modulation of cutaneous neurons was smaller than that of muscle-like neurons and was equally likely to be attenuation as potentiation. These widely varied patterns of sensitivity change, and their consistency within experimental sessions (Fig 8C), suggest that they are fine-tuned across muscles and receptors, perhaps “spotlighting” relevant information rather than the result of a more global effect.

CN neurons receive input both directly from peripheral receptors and by way of spinal interneurons in laminae 3-7. One study estimates that in the rat, between 30-40% of dorsal column afferents to CN are these second-order neurons (Giesler, Nahin, & Madsen, 1984; Loutit et al., 2021). Thus, gain modulation in CN might have a spinal origin. One study found cutaneous afferent input to cervical spinal interneurons to be consistently attenuated during active movements, while proprioceptive information was potentiated (Confais, Kim, Tomatsu, Takei, & Seki, 2017). That study differed from ours in the location of the receptive fields, ours focusing on neurons with proximal limb RFs, and the earlier study, the hand and distal arm. The discrepancy between our studies may result from the very different roles of distal cutaneous neurons for stereognosis and object interactions, and proximal arm neurons (both cutaneous and muscle) for control of reaching and a sense of limb position and movement. Importantly, our experiments could not distinguish between altered gamma drive, spinal modulation of spinal transmission, or descending inputs to CN as the source for the amplification of proprioception in our recordings.

### CN responses: A lens into gamma drive

The influence of gamma drive on spindle responses during active reaching movements is understood only qualitatively (Dimitriou & Edin, 2008; Proske & Gandevia, 2012). Our ability to record CN neurons during reaching may provide an indirect view of gamma modulation of spindle activity. In the passive condition, many spindle-receiving CN neurons reduced their firing for non-preferred directions, responses presumably associated with shortening of the muscle in their RF. However, these same neurons often did not have decreased rates during active movement in the same directions, suggesting that gamma drive may have prevented the spindles from falling silent. In fact, we often saw transient *increases* in the firing rate in these anti-preferred directions near movement onset (supplementary fig 7). These effects are consistent with increased gamma drive, though we cannot rule out other effects of descending modulatory input to the spinal cord or CN.

### Use of CN as a neural interface site for somatosensory replacement

With the increasing sophistication of efferent brain computer interfaces that can allow paralyzed patients to move (Collinger, Gaunt, & Schwartz, 2018; Hochberg et al., 2006; Lee et al., 2018), attention has swung to the complementary problem: restoring touch and proprioception to these patients by activating the somatosensory system electrically (Bensmaia & Miller, 2014; Flesher et al., 2016; Tabot et al., 2013). Somatosensory cortical stimulation has been used in both intact monkeys and paralyzed patients to elicit somatosensory percepts. Humans with electrode arrays implanted in the primary somatosensory cortex report strong, repeatable sensations from stimulation, including pressure, tingling, and vibration. However, proprioceptive-like percepts have been rare or absent (Collinger et al., 2018; Flesher et al., 2016; Lee et al., 2018). Likewise, targeted muscle reinnervation (TMR) and peripheral nerve stimulation have shown promise in restoring sensation in limb amputees (Horch, Meek, Taylor, & Hutchinson, 2011; Schiefer, Graczyk, Sidik, Tan, & Tyler, 2018; Tan et al., 2014), in part because the simpler coding and additional peripheral processing may simplify stimulus paradigms.

For spinal injury patients, the most peripheral site above the lesion is the CN, making it an appealing option to consider as a site of stimulation for sensory replacement (Loutit & Potas, 2020). We found a somatotopy across each array that was consistent across monkeys. Neurons were segregated both by modality (rostral and ventral subnuclei) and receptive field location, similar to earlier descriptions (Loutit et al., 2021). This somatotopic representation may allow for coherent proprioceptive percepts to be evoked via electrical stimulation. One drawback to CN as a site of proprioceptive replacement is the potential for damage to the dorsal columns or other medullary nuclei. While deafferentation is not a major concern for a person with a spinal cord injury who lacks sensation, CN lies close to medullary regions critical for homeostatic regulation, such as the dorsal respiratory group (Berger, 1977). Attempts to restore sensation in the medulla need to take care to minimize trauma to the surrounding tissue, perhaps with lower stiffness or non-penetrating electrodes.

## Acknowledgements

Thanks to Drs. Leah Krubitzer and Mary Baldwin for their histological expertise and helpful discussions.

---

**Sup. Fig 1:**
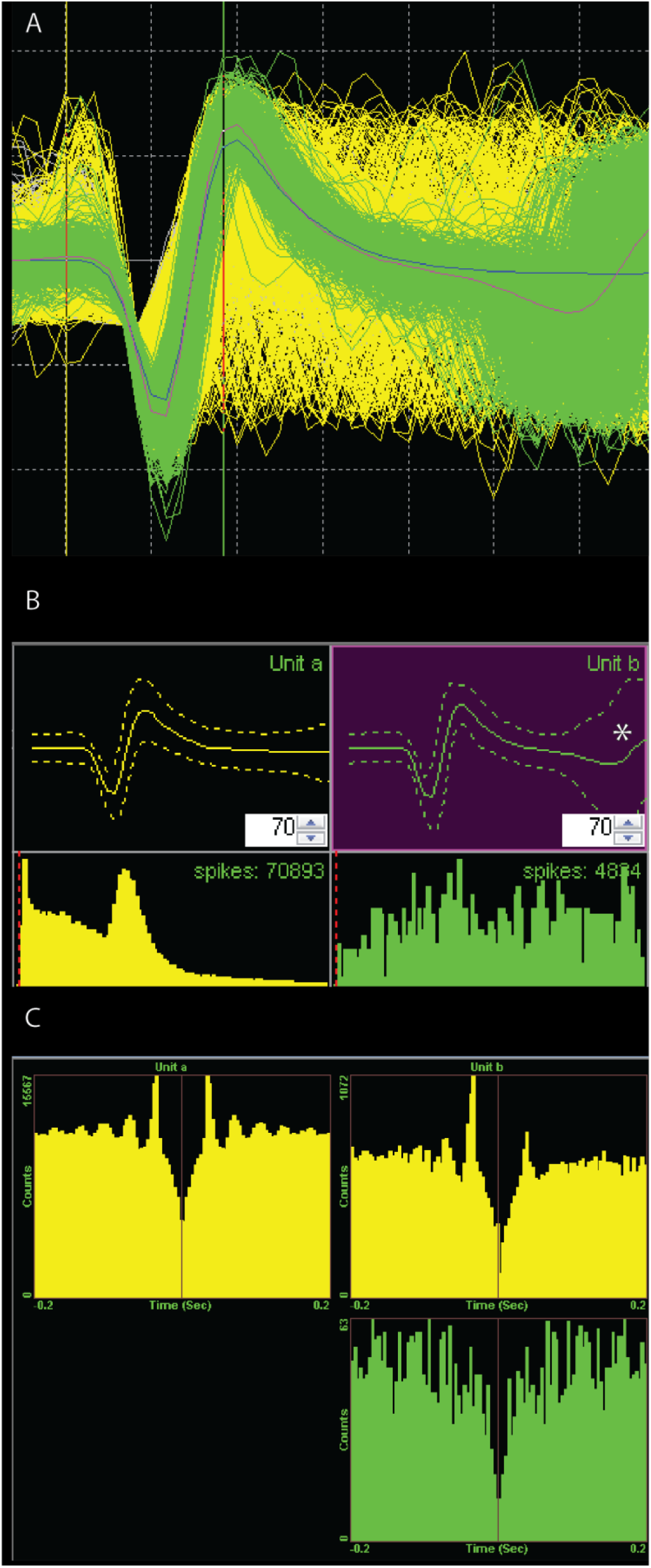
Example CN neuron that fires doublet spikes: A: Waveshape in Offline Sorter of single neuron in CN that fires in two separable modes, either a single spike or two spikes in a single window. Green neuron (sorted separately for visualization) has a large primary spike at the beginning of the 1.6 ms window and a second, temporally precise spike near the end of the same window (green deflections at the right side of the plot). Inter-spike interval (ISI) and cross correlation lags indicate that the yellow neuron is the same neuron without a double spike. B: Waveshapes (top row) and ISI histograms (bottom row) for the sorted waveshapes. Yellow neuron has a strong peak in its ISI, suggesting rhythmicity in its firing. CI at end of green waveshape expands as the second spike occurs, denoted with an asterisk. C: Cross correlogram between the two sorted waveshapes. Upper right quadrant depicts the relative timing between the yellow and green waveshape. Mass to the left of the red line indicates probability that the yellow spike precedes the green spike. This asymmetry was used to find groups of waveshapes that were likely generated by the same neuron.

**Sup. Fig. 2:**
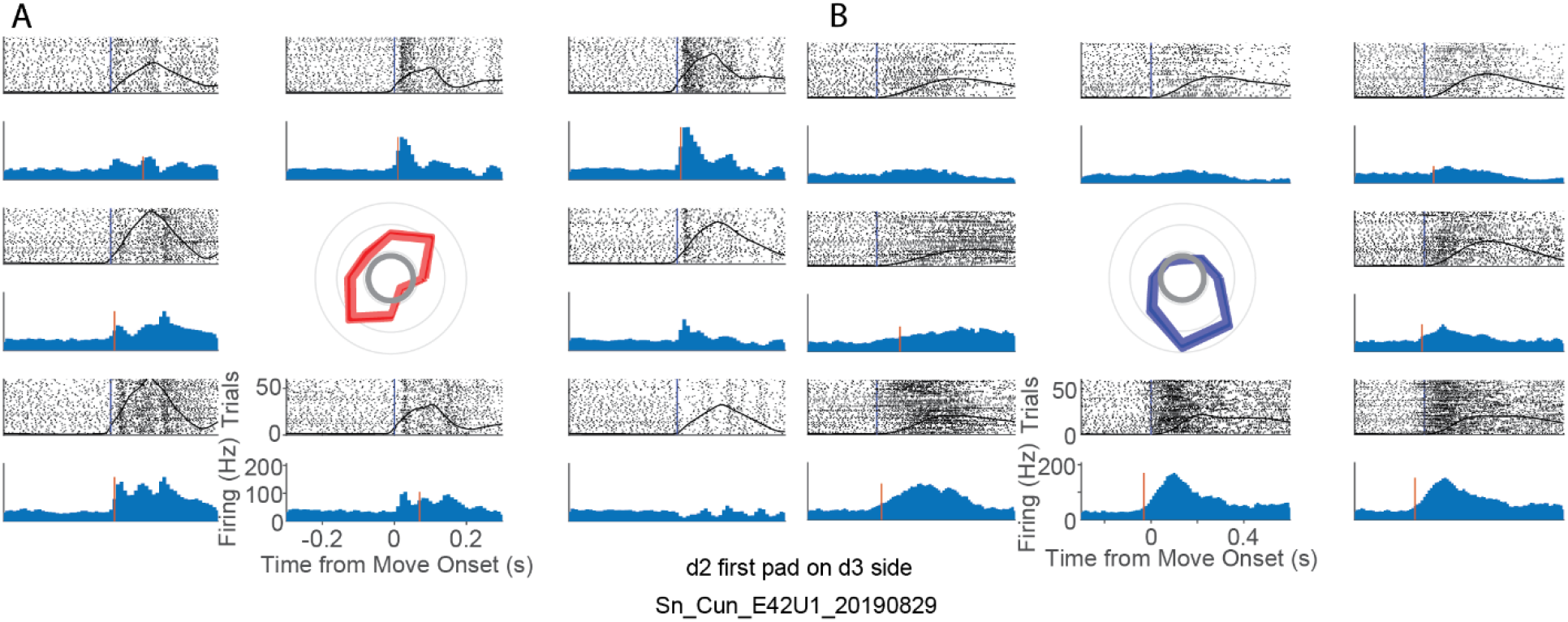
Example response of CN neuron that receives input from cutaneous receptors on the hand: A: Passive response of neuron that receives input from the d2 on the first pad, d3 side. Figure arranged with layout in Fig 5. Bimodal passive tuning curve (shown in center) was commonly observed for neurons with cutaneous receptive fields on the hand. B: Response of this neuron during actively generated reaching. We saw strong responses in both active and passive conditions but were not able to compare neural sensitivity across conditions due to the inconsistent relationship between hand kinematics and tactile responses.

**Sup. Table 1:** Movement-related firing rate statistics of CN neurons: Data from one recording session for each monkey are summarized here, with the date in parenthesis beneath the monkey. Neurons were included in the “cuneate” class if receptive fields were located on the arm or torso. Neurons not explicitly mapped on a given session were considered to be in CN if the electrode on which they were recorded was consistently mapped as CN on days in which it was tested. “Mapped” indicates how many neurons’ receptive fields were mapped across all a given monkey’s experimental sessions. “Spindle-receiving” indicates the number responsive to carefully-placed vibration, indicating a neuron potentially receiving muscle spindle inputs, while “muscle-like” included both these spindle-receiving and muscle-like neurons that did not respond to vibration (see further description in methods). “Cutaneous” neurons had an RF that responded to light touch, as described in the methods. “Proximal”, “elbow”, and “distal” neurons had receptive fields near the shoulder, elbow and wrist, respectively. Neurons with receptive fields that spanned a joint were counted for both segments; this was fairly common, minimally a result of input from a biarticular muscle or large a cutaneous field.

| Row | Combined | Bu<br>(20190329) | Sn<br>(20190829) | Cr<br>(20190418) |
| --- | --- | --- | --- | --- |
| Total | 212 | 106 | 59 | 47 |
| Cuneate | 126 | 37 | 56 | 33 |
| ActiveTuned | 109 | 28 | 50 | 31 |
| PassiveTuned | 76 | 14 | 31 | 31 |
| SinTunedAct | 111 | 26 | 53 | 32 |
| SinTunedPas | 67 | 12 | 28 | 27 |
| SinTunedBoth | 67 | 12 | 28 | 27 |
| Proximal | 27 | 9 | 6 | 12 |
| Elbow | 18 | 5 | 6 | 7 |
| Distal | 15 | 3 | 6 | 6 |
| Hand | 10 | 2 | 2 | 6 |
| Mapped | 82 | 24 | 26 | 32 |
| Cutaneous | 39 | 7 | 16 | 16 |
| Spindle-receiving | 40 | 17 | 9 | 14 |
| Muscle-like | 42 | 17 | 10 | 15 |

**Sup. Fig. 3:**
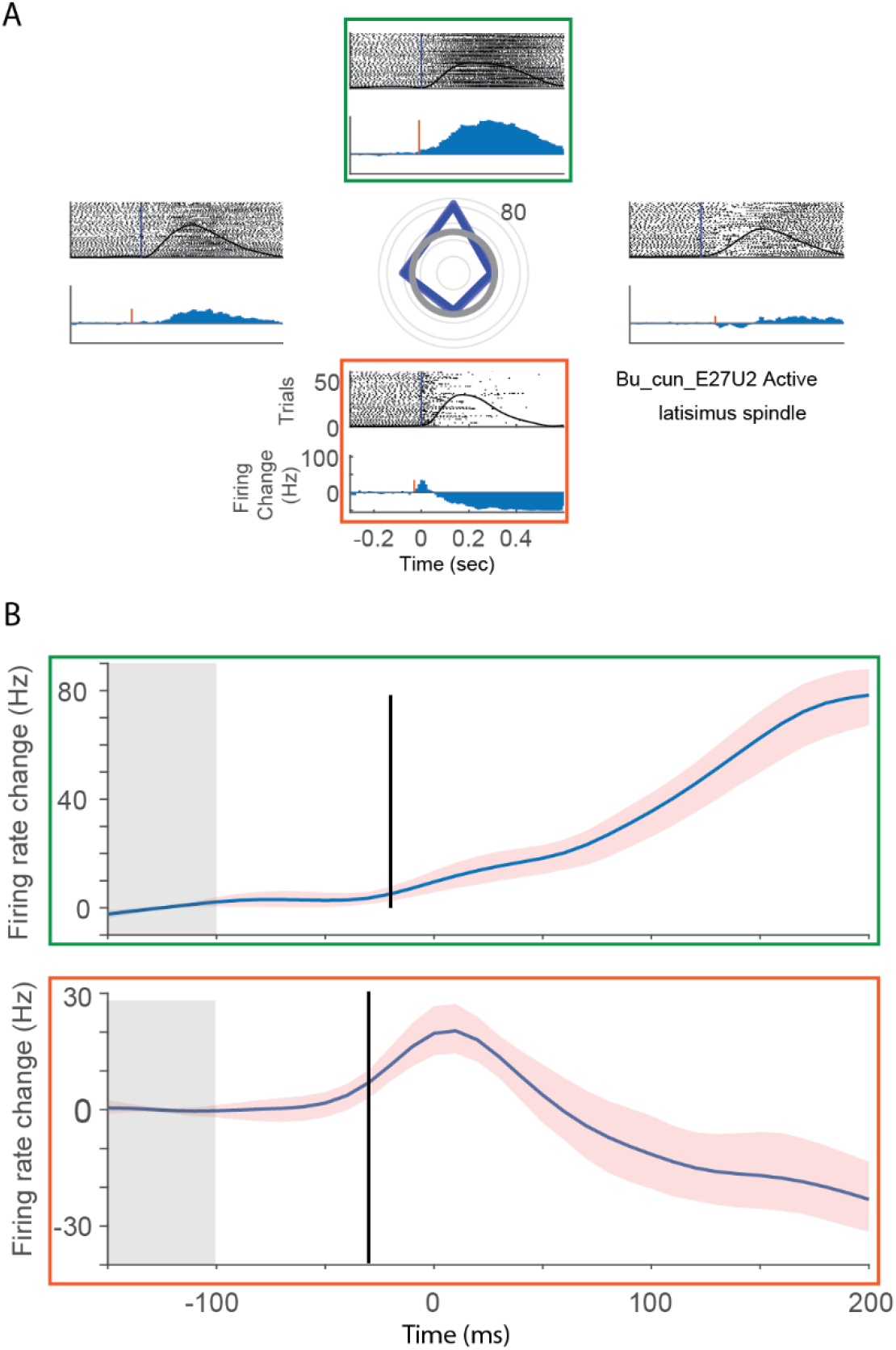
Illustration of changepoint analysis: A: Set of responses duplicated from Sup Fig 7A, active condition. B: Changepoint analysis of time of firing rate change near movement onset. Top, green plot-Average difference from baseline firing rate (shaded grey regions) over time for reaches away from the body (corresponding to green box in A). Shaded red area indicates the 95% confidence interval. A firing rate increase was detected ~20 ms prior to movement onset. Bottom - Same as top, except for reaches towards body (corresponding to orange box in A). Firing rate increase detected ~25 ms prior to movement onset.

**Sup. Fig. 4:**
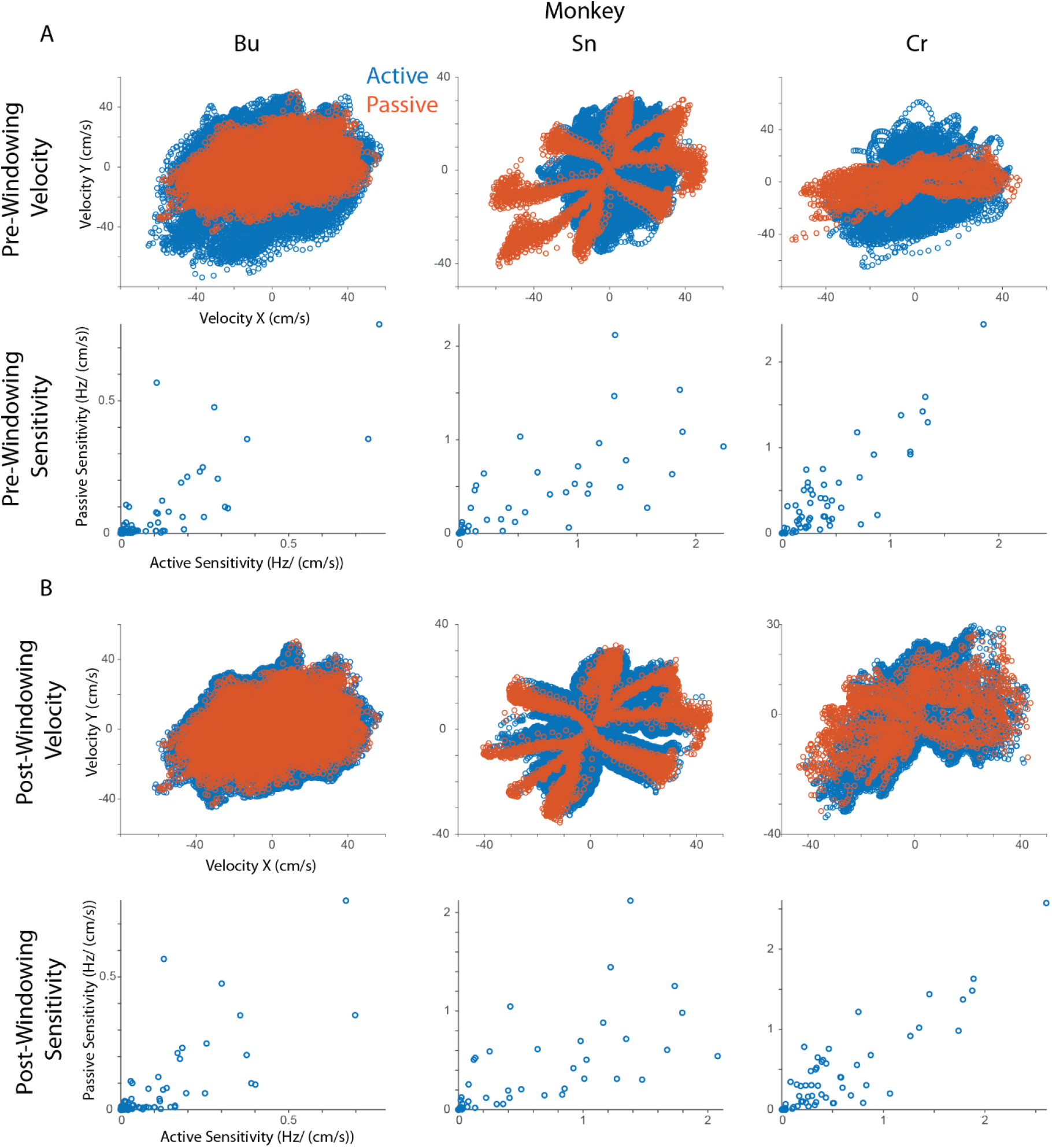
Mismatches in movement kinematics are corrected through a neighborhood data windowing technique: In CN monkeys, kinematics in the active (blue) and passive (orange) conditions differ due to the anisotropy of the arm impedance and musculature. A: First Row - Velocity space in the 400 ms analysis window after movement onset during reach (blue) and the 130 ms window after bump onset (orange). Second row: Sensitivities in the passive condition plotted against the active condition for each monkey. B: Same as top row of A, except nonoverlapping velocity data has been removed. Bottom row: Same as second row after windowing. The computed sensitivities do not change substantially indicating that the models fit to compute sensitivity seem robust to the data windowing process.

**Sup. Fig. 5:**
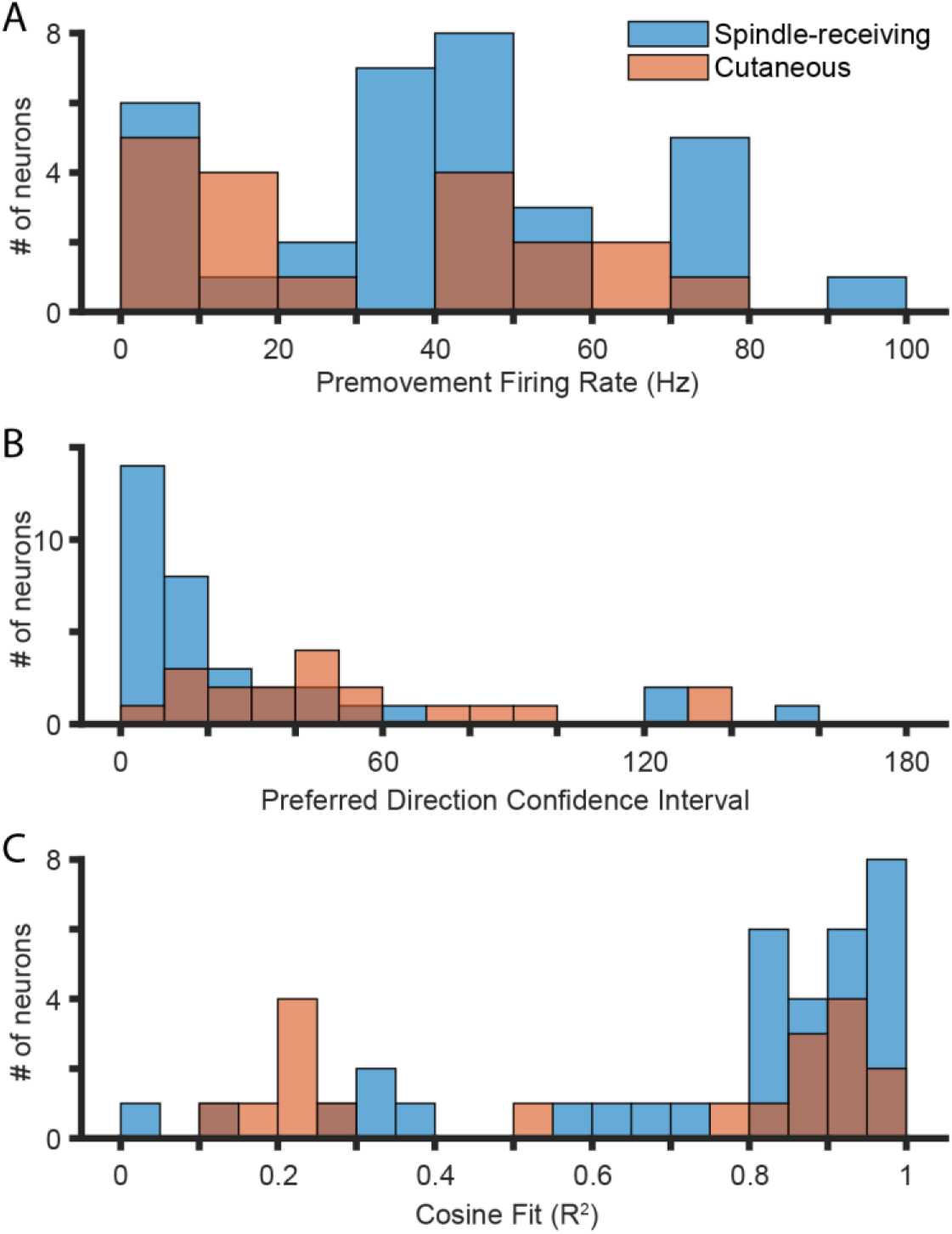
Firing properties of spindle-receiving and cutaneous CN neurons: Plot includes all CN neurons that do not have RFs on the distal arm. A: Histogram of mean pre-movement firing rates for spindle-receiving CN neurons (blue) and cutaneous neurons (pink). Spindle-receiving neurons often had higher baseline firing rates than cutaneous neurons, there was no statistically significant difference (KS-test, p≈0.13). B: Histogram of PD confidence interval for spindle and cutaneous CN neurons. Spindle-receiving neurons typically had tighter confidence intervals than cutaneous neurons (KS test, p ≈ 0.006). C: Sinusoidal goodness-of-fit (R^2^ of cosine tuning model) for these neurons. There was no difference between the modality types (KS test, p ≈0.24).

**Sup. Fig. 6:**
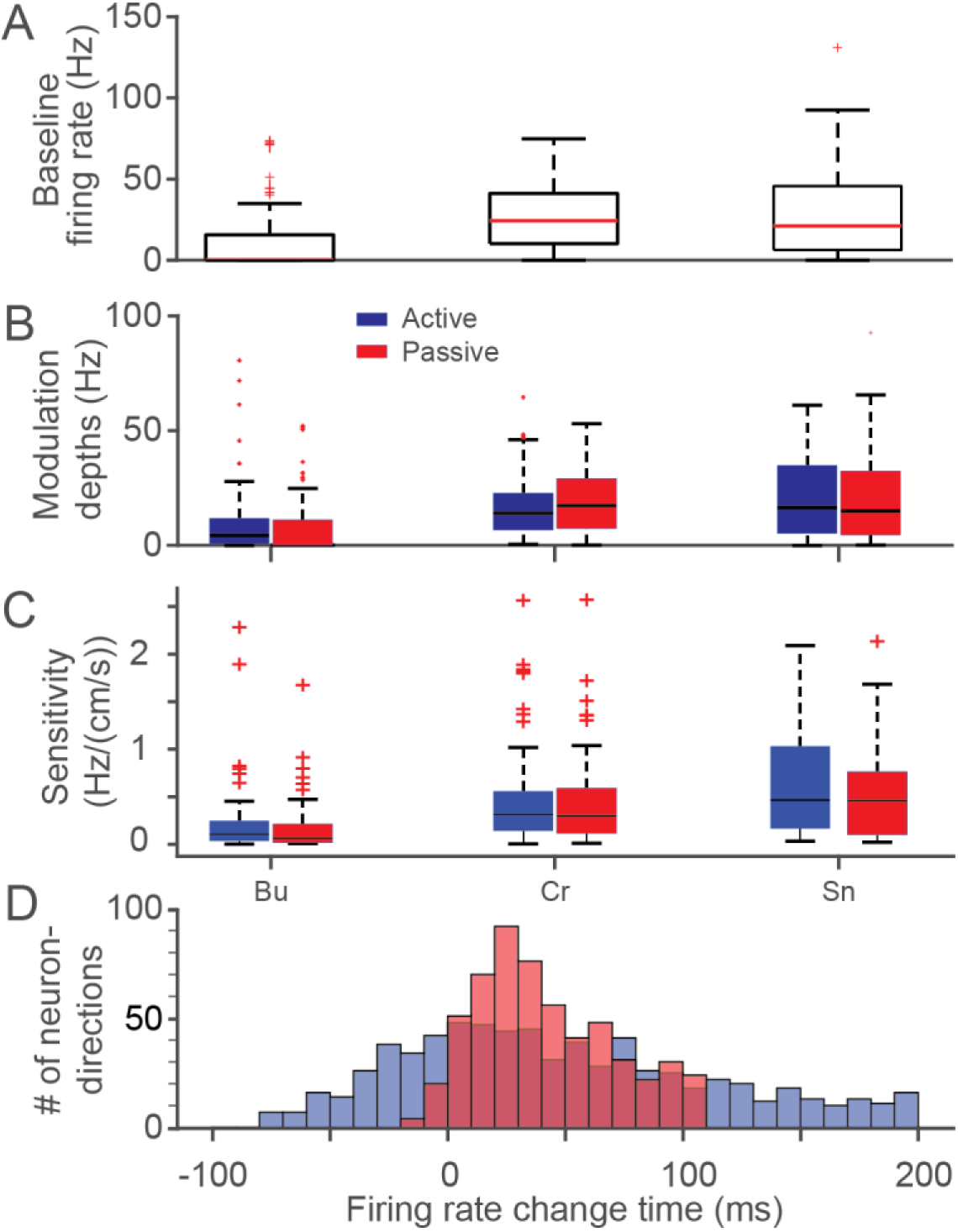
Response statistics of CN neurons during active and passive movements: A: Box and whisker plot of baseline (100 to 50 ms before movement onset) firing rate of CN neurons from three monkeys (Bu, Cr, Sn). Red line indicates the median, and top and bottom of box indicate the 75% and 25% quantiles respectively. Red crosses indicate outlier neurons. B: Spatial tuning curve depths in active (blue) and passive (red) conditions. C: Sensitivity of CN neurons to hand speed. D: Histogram of response latency relative to movement onset, marginalized across monkeys.

**Sup. Fig. 7:**
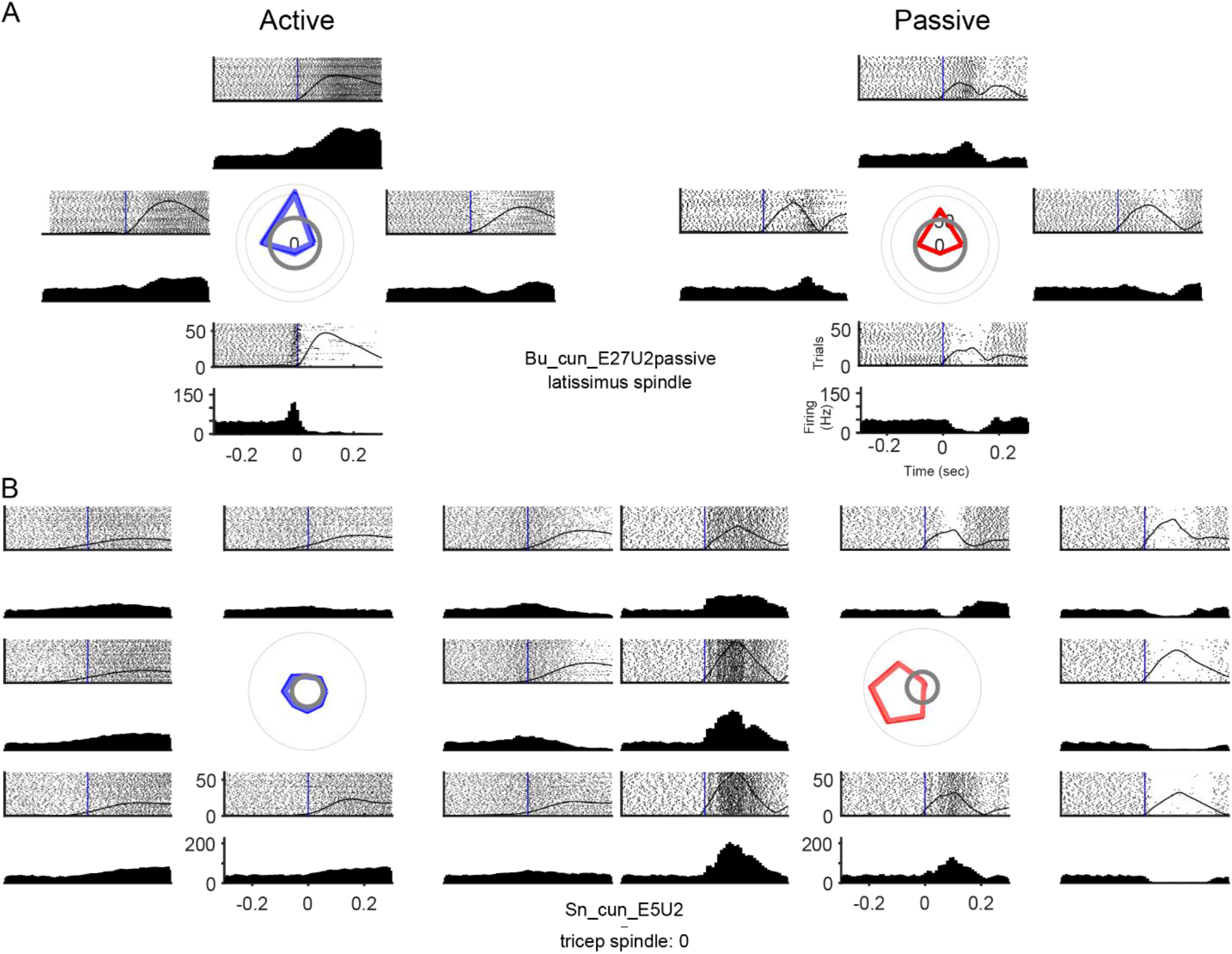
Example spindle-receiving neurons with pre-movement firing rate increases in the anti-preferred direction: A: A CN neuron that appeared to receive muscle spindle inputs from the latissimus. Preferred direction points away from the body in both active (left column, blue) and passive (right column, red) conditions. Rasters and trial averaged firing rates arranged radially as in Fig 4. There is a sharp increase in the anti-preferred direction (in this case, towards the body) in the active condition. B: Same as A, except showing a neuron with inputs from triceps muscle spindles. The PD points to the left in both conditions, but the firing rate in the active case increases in the anti-preferred direction (to the right) prior to movement. This unit is also one of the rare examples in which the active response in a spindle-receiving neuron was substantially weaker than the passive response.

**Sup. Fig. 8:**
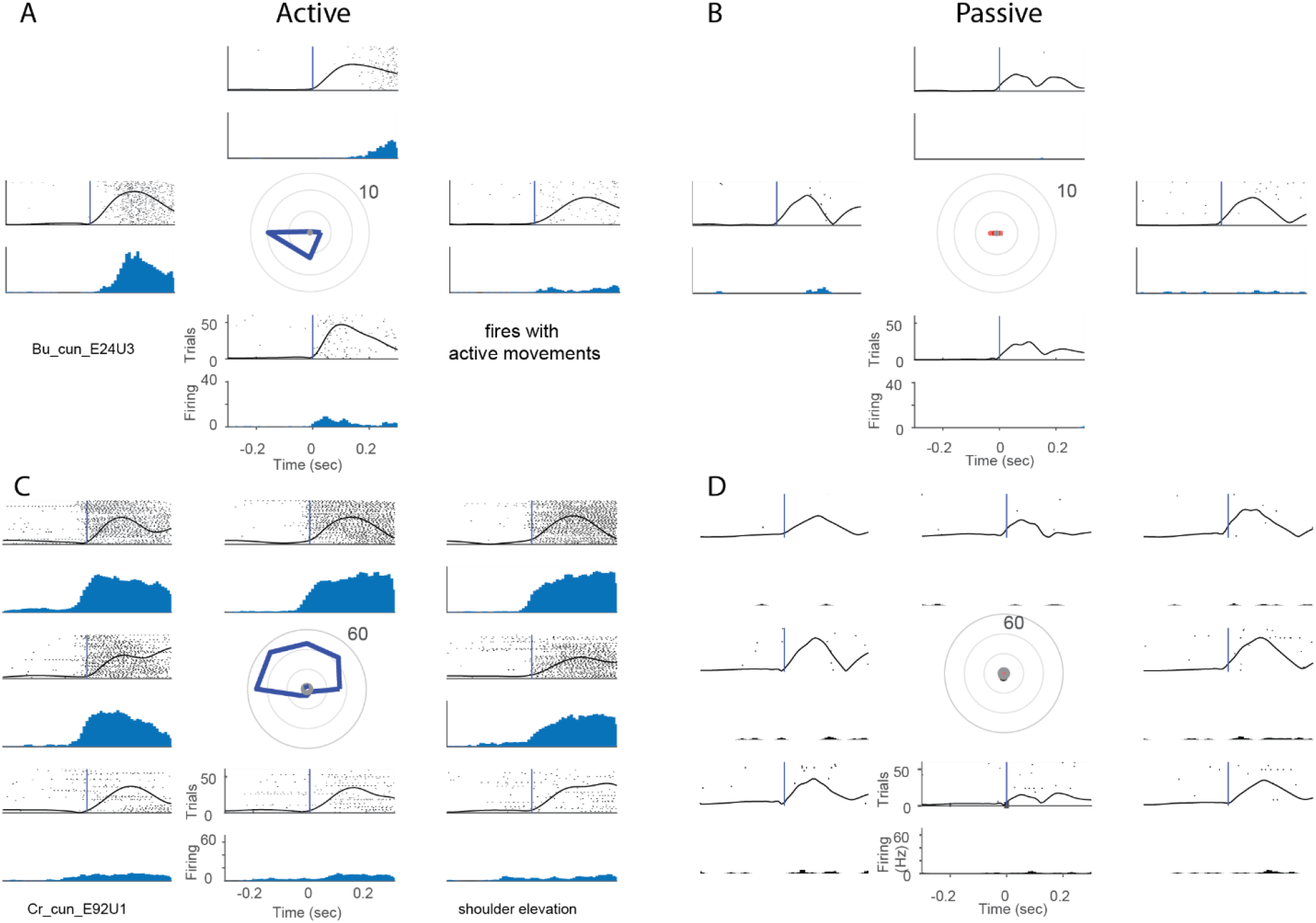
Example proprioceptive neurons strongly activated in active, but not passive conditions: A: Example neuron that responded when the monkey made active movements, but not during passive displacement. High firing rate occurred in leftwards reaches. B: Same neuron as A was unresponsive during passive perturbation. C: A CN neuron that responded to shoulder elevation and when reaching away from the body. D: This neuron was also unresponsive during passive perturbation.

**Sup. Fig. 9:**
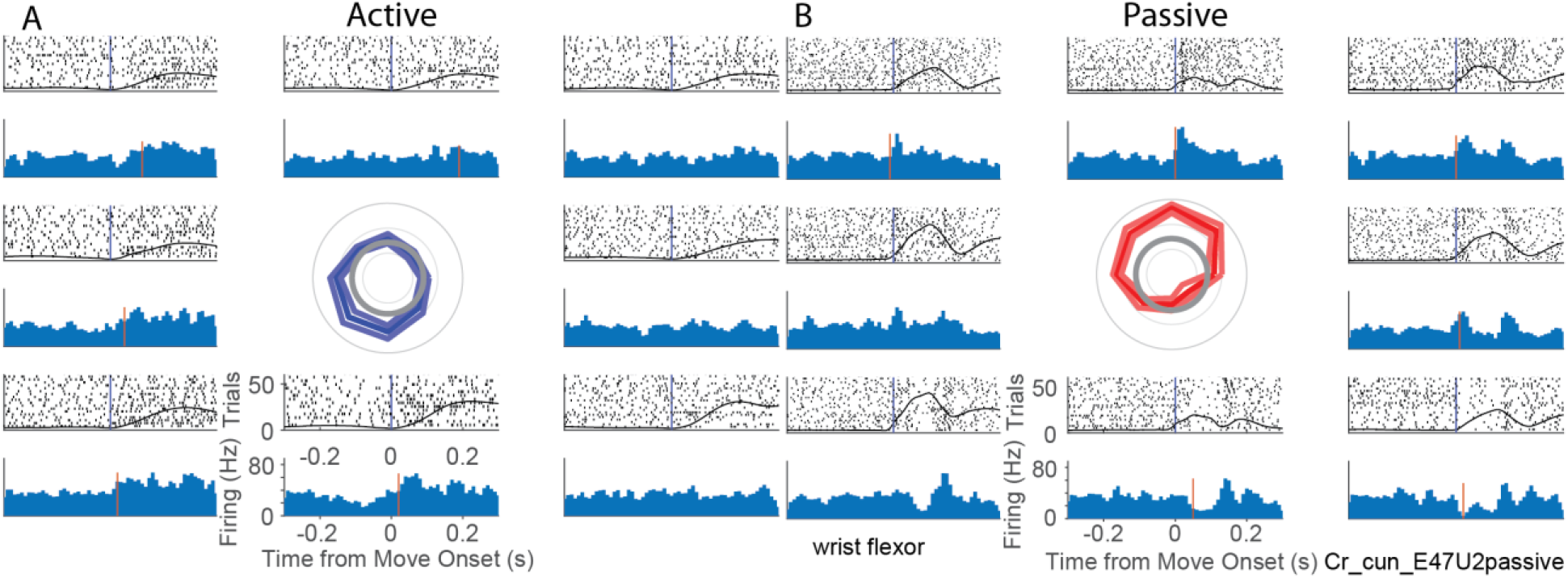
Example active and passive responses of neuron that appeared to receive inputs from wrist flexor muscle spindles: A: Responses of a neuron during active reach. Highest firing rates in the active condition were during reaches toward the body. B: Same neuron as A. Strongest passive responses are evident at bump onset for bumps away from the body and in most other directions at ~150 ms after bump onset. The complex differences in dynamics of the distal arm during reach and perturbation cause the kinematic PDs to point in different directions.

**Sup. Fig. 10:**
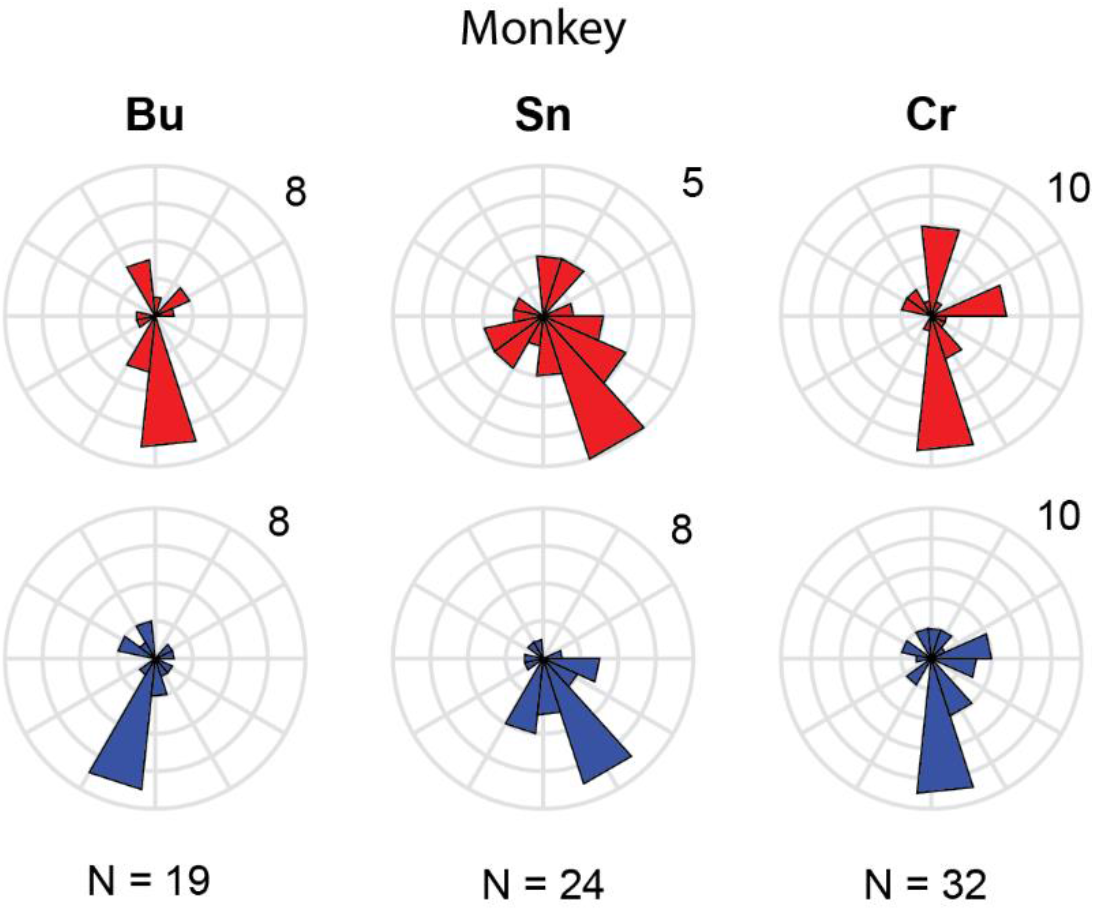
PD distributions for Individual monkeys: Neurons included in this figure correspond to the combined distributions in Fig 6, and include those that were sinusoidally tuned in both active and passive conditions, from CN regions of the array that appeared to receive inputs from the upper trunk, shoulder or proximal arm. Passive PD distributions (red) are on the top row, active PD distributions (blue) are on the bottom row.

**Sup. Fig. 11:**
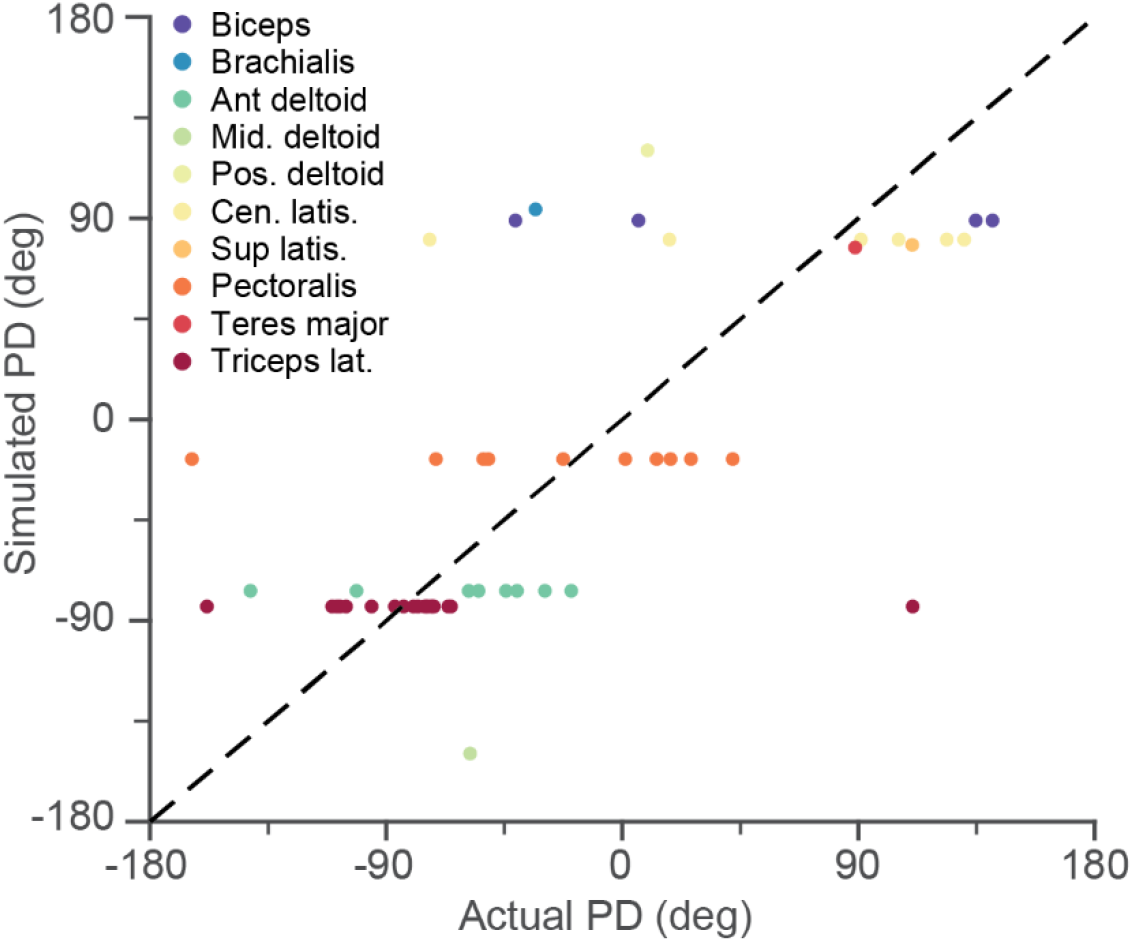
Comparison of actual and simulated PDs: Scatter plot relating the empirically computed active PD for a given neuron to the expected PD if it received input only from its single dominant apparent spindle input. Each point represents a single neuron, color-coded by its muscle RF.

## References

Andersen, P., Eccles, J. C., Oshima, T., & Schmidt, R. F. (1964). Mechanisms of synaptic transmission in the Cuneate Nucleus. Journal of Neurophysiology, 27(6), 1096–1116. Retrieved from http://www.ncbi.nlm.nih.gov/pubmed/14223972

Andersen, P., Eccles, J. C., Schmidt, R. F., & Yokota, T. (1964). Identification of relay cells and interneurons in the Cuneate Nucleus. Journal of Neurophysiology, 27(6), 1080–1095. https://doi.org/10.1152/jn.1964.27.6.1080

Balaram, P., Young, N. A., & Kaas, J. H. (2014). Histological features of layers and sublayers in cortical visual areas V1 and V2 of chimpanzees, macaque monkeys, and humans. Eye and Brain, 6(Suppl 1), 5–18. https://doi.org/10.2147/EB.S51814

Banks, R., & Stacey, M. (1988). Quantitative Studies on Mammalian Muscle Spindles and their Sensory Innervation. In Mechanoreceptors (pp. 263–269). Springer US. https://doi.org/10.1007/978-1-4899-0812-4_49

Bengtsson, F., Brasselet, R., Johansson, R. S., Arleo, A., & Jörntell, H. (2013). Integration of Sensory Quanta in Cuneate Nucleus Neurons In Vivo. PLoS ONE, 8(2). https://doi.org/10.1371/journal.pone.0056630

Bensmaia, S. J., & Miller, L. E. (2014, May). Restoring sensorimotor function through intracortical interfaces: Progress and looming challenges. Nature Reviews Neuroscience. Nature Publishing Group. https://doi.org/10.1038/nrn3724

Berger, A. J. (1977). Dorsal respiratory group neurons in the medulla of cat: Spinal projections, responses to lung inflation and superior laryngeal nerve stimulation. Brain Research, 135(2), 231–254. https://doi.org/10.1016/0006-8993(77)91028-9

Chan, S. S., & Moran, D. W. (2006). Computational model of a primate arm: from hand position to joint angles, joint torques and muscle forces. J. Neural Eng, 3, 327–337. https://doi.org/10.1088/1741-2560/3/4/010

Chowdhury, R. H., Glaser, J. I., & Miller, L. E. (2020). Area 2 of primary somatosensory cortex encodes kinematics of the whole arm. ELife, 9. https://doi.org/10.7554/eLife.48198

Cohen, L. G., & Starr, A. (1987). Localization, Timing And Specificity Of Gating Of Somatosensory Evoked Potentials During Active Movement In Man (Vol. 110). Retrieved from https://academic.oup.com/brain/article-abstract/110/2/451/467280

Collinger, J. L., Gaunt, R. A., & Schwartz, A. B. (2018). Progress towards restoring upper limb movement and sensation through intracortical brain-computer interfaces. Current Opinion in Biomedical Engineering, 8, 84–92. https://doi.org/10.1016/J.COBME.2018.11.005

Collins, D. F., Refshauge, K. M., Todd, G., & Gandevia, S. C. (2005). Cutaneous receptors contribute to kinesthesia at the index finger, elbow, and knee. Journal of Neurophysiology, 94(3), 1699–1706. https://doi.org/10.1152/jn.00191.2005

Confais, J., Kim, G., Tomatsu, S., Takei, T., & Seki, K. (2017). Nerve-specific input modulation to spinal neurons during a motor task in the monkey. Journal of Neuroscience, 37(10), 2612–2626. https://doi.org/10.1523/JNEUROSCI.2561-16.2017

Delp, S. L., Anderson, F. C., Arnold, A. S., Loan, P., Habib, A., John, C. T., … Thelen, D. G. (2007). OpenSim: Open-source software to create and analyze dynamic simulations of movement. IEEE Transactions on Biomedical Engineering, 54(11), 1940–1950. https://doi.org/10.1109/TBME.2007.901024

Dimitriou, M. (2014). Human muscle spindle sensitivity reflects the balance of activity between antagonistic muscles. The Journal of Neuroscience: The Official Journal of the Society for Neuroscience, 34(41), 13644–13655. https://doi.org/10.1523/JNEUROSCI.2611-14.2014

Dimitriou, M., & Edin, B. B. (2008). Discharges in human muscle spindle afferents during a key-pressing task. Journal of Physiology, 586(22), 5455–5470. https://doi.org/10.1113/jphysiol.2008.160036

Edin, B. B. (1992). Quantitative analysis of static strain sensitivity in human mechanoreceptors from hairy skin. Journal of Neurophysiology, 67(5), 1105–1113. https://doi.org/10.1152/jn.1992.67.5.1105

Edin, B. B., & Johansson, N. (1995). Skin strain patterns provide kinaesthetic information to the human central nervous system. The Journal of Physiology, 487(1), 243–251. https://doi.org/10.1113/jphysiol.1995.sp020875

Fallon, J. B., & Macefield, V. G. (2007). Vibration sensitivity of human muscle spindles and Golgi tendon organs. Muscle & Nerve, 36(1), 21–29. https://doi.org/10.1002/mus.20796

Flesher, S. N., Collinger, J. L., Foldes, S. T., Weiss, J. M., Downey, J. E., Tyler-Kabara, E. C., … Gaunt, R. A. (2016). Intracortical microstimulation of human somatosensory cortex. Science Translational Medicine, 8(361), 361ra141–361ra141. https://doi.org/10.1126/scitranslmed.aaf8083

Georgopoulos, A. P., Kalaska, J. F., Caminiti, R., & Massey, J. T. (1982). On the relations between the direction of two-dimensional arm movements and cell discharge in primate motor cortex. The Journal of Neuroscience : The Official Journal of the Society for Neuroscience, 2(11), 1527–1537. Retrieved from http://www.ncbi.nlm.nih.gov/pubmed/7143039

Ghez, C., & Pisa, M. (1972). Inhibition of afferent transmission in cuneate nucleus during voluntary movement in the cat. Brain Research, 40(1), 145–155. Retrieved from http://www.ncbi.nlm.nih.gov/pubmed/4338259

Giesler, G. J., Nahin, R. L., & Madsen, A. M. (1984). Postsynaptic dorsal column pathway of the rat. I. Anatomical studies. Journal of Neurophysiology, 51(2), 260–275. https://doi.org/10.1152/jn.1984.51.2.260

He, Q., Suresh, A., Versteeg, C., Rosenow, J. M., & Bensmaia, S. J. (2019). Movement gating of cutaneous signals in the cuneate nucleus. Presented at Society for Neuroscience 2019.

Hochberg, L. R., Serruya, M. D., Friehs, G. M., Mukand, J. A., Saleh, M., Caplan, A. H., … Donoghue, J. P. (2006). Neuronal ensemble control of prosthetic devices by a human with tetraplegia. Nature, 442(7099), 164–171. https://doi.org/10.1038/nature04970

Horch, K., Meek, S., Taylor, T. G., & Hutchinson, D. T. (2011). Object discrimination with an artificial hand using electrical stimulation of peripheral tactile and proprioceptive pathways with intrafascicular electrodes. IEEE Transactions on Neural Systems and Rehabilitation Engineering, 19(5), 483–489. https://doi.org/10.1109/TNSRE.2011.2162635

Houk, J. C., Rymer, W. Z., & Crago, P. E. (1981). Dependence of dynamic response of spindle receptors on muscle length and velocity. Journal of Neurophysiology, 46(1), 143–166. https://doi.org/10.1152/jn.1981.46.1.143

Hummelsheim, H., & Wiesendanger, M. (1985). Neuronal responses of medullary relay cells to controlled stretches of forearm muscles in the monkey. Neuroscience, 16(4), 989–996. https://doi.org/10.1016/0306-4522(85)90111-3

Juravle, G., Binsted, G., & Spence, C. (2017). Tactile suppression in goal-directed movement. Psychonomic Bulletin and Review, 24(4), 1060–1076. https://doi.org/10.3758/s13423-016-1203-6

Lee, B., Kramer, D., Salas, M. A., Kellis, S., Brown, D., Dobreva, T., … Andersen, R. A. (2018). Engineering artificial somatosensation through cortical stimulation in humans. Frontiers in Systems Neuroscience, 12. https://doi.org/10.3389/fnsys.2018.00024

Lillicrap, T. P., & Scott, S. H. (2013). Preference Distributions of Primary Motor Cortex Neurons Reflect Control Solutions Optimized for Limb Biomechanics. Neuron, 77(1), 168–179. https://doi.org/10.1016/J.NEURON.2012.10.041

London, B. M., & Miller, L. E. (2013). Responses of somatosensory area 2 neurons to actively and passively generated limb movements. Journal of Neurophysiology, 109(6), 1505–1513. https://doi.org/10.1152/jn.00372.2012

Loutit, A. J., & Potas, J. R. (2020). Restoring Somatosensation: Advantages and Current Limitations of Targeting the Brainstem Dorsal Column Nuclei Complex. Frontiers in Neuroscience, 14, 156. https://doi.org/10.3389/fnins.2020.00156

Loutit, A. J., Vickery, R. M., & Potas, J. R. (2021). Functional organization and connectivity of the dorsal column nuclei complex reveals a sensorimotor integration and distribution hub. Journal of Comparative Neurology, 529(1), 187–220. https://doi.org/10.1002/cne.24942

Mathis, A., Mamidanna, P., Cury, K. M., Abe, T., Murthy, V. N., Mathis, M. W., & Bethge, M. (2018). DeepLabCut: markerless pose estimation of user-defined body parts with deep learning. Nature Neuroscience, 21(9), 1281–1289. https://doi.org/10.1038/s41593-018-0209-y

Palmeri, A., Bellomo, M., Giuffrida, R., & Sapienza, S. (1999). Motor cortex modulation of exteroceptive information at bulbar and thalamic lemniscal relays in the cat. Neuroscience, 88(1), 135–150. Retrieved from http://www.ncbi.nlm.nih.gov/pubmed/10051195

Proske, U., & Gandevia, S. C. (2012). The Proprioceptive Senses: Their Roles in Signaling Body Shape, Body Position and Movement, and Muscle Force. Physiological Reviews, 92(4), 1651–1697. https://doi.org/10.1152/physrev.00048.2011

Prud’homme, M. J., & Kalaska, J. F. (1994). Proprioceptive activity in primate primary somatosensory cortex during active arm reaching movements. Journal of Neurophysiology, 72(5), 2280–2301. Retrieved from http://www.ncbi.nlm.nih.gov/pubmed/7884459

Rushton, D. N., Rothwell, J. C., & Craggs, M. D. (1981). Gating Of Somatosensory Evoked Potentials During Different Kinds Of Movement In Man. Brain (Vol. 104). Retrieved from https://academic.oup.com/brain/article-abstract/104/3/465/296781

Sainburg, R. L., Ghilardi, M. F., Poizner, H., & Ghez, C. (1995). Control of limb dynamics in normal subjects and patients without proprioception. Journal of Neurophysiology, 73(2), 820–835. https://doi.org/10.1152/jn.1995.73.2.820

Sandbrink, K., Mamidanna, P., Michaelis, C., Mathis, M. W., Bethge, M., & Mathis, A. (2020). Task-driven hierarchical deep neural network models of the proprioceptive pathway. https://doi.org/10.1101/2020.05.06.081372

Schiefer, M. A., Graczyk, E. L., Sidik, S. M., Tan, D. W., & Tyler, D. J. (2018). Artificial tactile and proprioceptive feedback improves performance and confidence on object identification tasks. PLOS ONE, 13(12), e0207659. https://doi.org/10.1371/journal.pone.0207659

Schmidt, R. F., Schady, W. J. L., & Torebjörk, H. E. (1990). Gating of tactile input from the hand - I. Effects of finger movement. Experimental Brain Research, 79(1), 97–102. https://doi.org/10.1007/BF00228877

Stevenson, I. H., Cherian, A., London, B. M., Sachs, N. A., Lindberg, E., Reimer, J., … Kording, K. P. (2011). Statistical assessment of the stability of neural movement representations. Journal of Neurophysiology, 106(2), 764–774. https://doi.org/10.1152/jn.00626.2010

Suresh, A. K., Winberry, J. E., Versteeg, C., Chowdhury, R., Tomlinson, T., Rosenow, J. M., … Bensmaia, S. J. (2017). Methodological considerations for a chronic neural interface with the cuneate nucleus of macaques. Journal of Neurophysiology, 118(6), 3271–3281. https://doi.org/10.1152/jn.00436.2017

Tabot, G. A., Dammann, J. F., Berg, J. A., Tenore, F. V., Boback, J. L., Vogelstein, R. J., & Bensmaia, S. J. (2013). Restoring the sense of touch with a prosthetic hand through a brain interface. Proceedings of the National Academy of Sciences of the United States of America, 110(45), 18279–18284. https://doi.org/10.1073/pnas.1221113110

Tan, D. W., Schiefer, M. A., Keith, M. W., Anderson, J. R., Tyler, J., & Tyler, D. J. (2014). A neural interface provides long-term stable natural touch perception. Science Translational Medicine, 6(257). https://doi.org/10.1126/scitranslmed.3008669

Williams, S. R., & Chapman, C. E. (2000). Time course and magnitude of movement-related gating of tactile detection in humans. II. Effects of stimulus intensity. Journal of Neurophysiology, 84(2), 863–875. https://doi.org/10.1152/jn.2000.84.2.863

Williams, S. R., & Chapman, C. E. (2002). Time course and magnitude of movement-related gating of tactile detection in humans. III. Effect of motor tasks. Journal of Neurophysiology, 88(4), 1968–1979. https://doi.org/10.1152/jn.2002.88.4.1968

Williams, S. R., Shenasa, J., & Chapman, C. E. (1998). Time Course and Magnitude of Movement-Related Gating of Tactile Detection in Humans. I. Importance of Stimulus Location. Journal of Neurophysiology, 79(2), 947–963. https://doi.org/10.1152/jn.1998.79.2.947

Witham, C. L., & Baker, S. N. (2011). Modulation and transmission of peripheral inputs in monkey cuneate and external cuneate nuclei. Journal of Neurophysiology, 106(5), 2764–2775. https://doi.org/10.1152/jn.00449.2011

